# Synergistic Effects of the DRD2/3 Antagonist ONC201 and Radiation in Glioblastoma

**DOI:** 10.1101/2020.07.23.218446

**Authors:** Ling He, Kruttika Bhat, Angeliki Ioannidis, Le Zhang, Nhan T. Nguyen, Joshua E. Allen, Phioanh Leia Nghiemphu, Timothy F. Cloughesy, Linda M. Liau, Harley I. Kornblum, Frank Pajonk

**Author notes:** **Authorship**: LH, KB, AI, LZ performed the experiments and collected the data. LML and HIK provided the patient-derived specimen. JEA provided ONC201, FP designed the experiments, FP conceived of the study, LH and FP analyzed the data and wrote the manuscript. All authors edited and approved of the final version of the manuscript. **Correspondence address**: Frank Pajonk, MD, PhD, Department of Radiation Oncology, David Geffen School of Medicine at UCLA, 10833 Le Conte Ave, Los Angeles, CA 90095-1714.

## Abstract

**Background:** Glioblastoma (GBM) is the deadliest of all brain cancers in adults. The current standard-of-care is surgery followed by radiotherapy and temozolomide, leading to a median survival time of only 15 months. GBM are organized hierarchically with a small number of glioma-initiating cells, responsible for therapy resistance and tumor recurrence, suggesting that targeting glioma-initiating cells could improve treatment response. ONC201 is a first-in-class anti-tumor agent with clinical efficacy in some forms of high-grade gliomas. Here we test its efficacy against GBM in combination with radiation.

**Methods:** Using patient-derived GBM lines and mouse models of GBM we test the effects of radiation and ONC201 on GBM self - renewal *in vitro* and survival *in vivo*. A possible resistance mechanism is investigated using RNA-Sequencing.

**Results:** Treatment of GBM cells with ONC201 reduced self-renewal, clonogenicity and cell viability *in vitro*. ONC201 exhibited anti-tumor effects on radioresistant GBM cells indicated by reduced self-renewal in secondary and tertiary glioma spheres. Combined treatment with ONC201 and radiation prolonged survival in syngeneic and patient-derived orthotopic xenograft mouse models of GBM. Subsequent transcriptome analyses after combined treatment revealed shifts in gene expression signatures related to quiescent GBM populations, GBM plasticity, and GBM stem cells.

**Conclusions:** Our findings suggest that combined treatment with the DRD2/3 antagonist ONC201 and radiation improves the efficacy of radiation against GBM *in vitro* and *in vivo* through suppression of GICs without increasing toxicity in mouse models of GBM. A clinical assessment of this novel combination therapy against GBM is further warranted.

**Key points:**

- Combined treatment of ONC201 and radiation exhibit anti-tumor effects on cells from primary and recurrent GBM
- Combined treatment significantly prolongs survival in vivo
- Combined treatment potentially targets the quiescent GBM cell population

**Importance of the Study:** The survival rates for patients with GBM are unacceptably low and novel treatment approaches are needed. This study provides evidence that a combination of radiation and the dopamine receptor antagonist ONC201 significantly prolongs survival in mouse models of glioma.

## Introduction

GBM is the deadliest of all brain cancers in adults and all patients ultimately succumb to the tumor. The standard-of-care involves surgical removal of the bulk tumor followed by radiotherapy and temozolomide treatment^1–4^. Reasons for treatment failure include the spread of tumor cells into the normal parenchyma, far beyond the detectable tumor, radio- and chemo-therapy resistance of the tumor cells and the intratumoral heterogeneity and plasticity of GBM^5–9^. Past attempts to improve the outcome of these patients using classical chemotherapeutic drugs or immunotherapy have been hampered by the inability of many drugs or biologics to cross the blood brain barrier (BBB)^10–12^.

ONC201 is the first member of a novel class of anti-cancer small molecules called imipridones, originally discovered by screening with the luciferase-reporter system for p53-independent inducers of the immuno-surveillance cytokine TNF-related apoptosis-inducing ligand (TRAIL) and tumor cell death in human colorectal cancer cells^13,14^. ONC201 has been found to specifically bind to and inhibit the dopamine receptor D2-like receptors DRD2 and DRD3 and acts as an allosteric agonist of the mitochondrial protease caseinolytic protease P (ClpP)^15^. This results in dual inactivation of Akt/ERK signaling that drives Foxo3a-mediated TRAIL gene induction to promote pro-apoptotic and anti-proliferative effects in tumor cells, but not in normal cells^16–20^. With p53-independency, low toxicity and excellent penetration of the BBB, ONC201 is currently being evaluated in multiple clinical trials for select advanced malignancies, including myeloma, leukemia, lymphoma, endometrial cancer, high grade glioma, and other solid tumors^21–28^.

Emerging data suggest that a hierarchical organization within glioblastoma plays a crucial role in tumor development, therapy resistance and tumor recurrence^29–32^ and interactions between tumor cells and the microenvironment are involved in tumor progression and the invasive nature of GBM^33,34^. Strategies to eradicate glioma-initiating cells at the apex of this hierarchy could be a valid approach for developing novel therapeutic approaches or modifying existing treatment regimen. Furthermore, GBM comprises of both a fast-dividing population and a relatively quiescent population, while conventional chemo and radiation therapies largely only target the proliferative population^35–38^. Hence, mobilizing the quiescent cell population into proliferation to further sensitize them to existing therapies could be an approach to improve GBM treatment outcome.

In this study we demonstrate that the first-in-class compound of imipridones, ONC201, in combination with radiation has anti-tumor efficacy against GBM *in vitro* and prolongs overall survival in mouse models of GBM *in vivo*. Transcriptome analyses revealed shifts in gene expression signatures in response to radiation and ONC201, indicating effects of the combination treatment on quiescent GBM cells and their ability to interact with the extracellular matrix (ECM).

## Material and Methods

### Reagents

ONC201 was kindly provided by Oncoceutics, Inc. (Philadelphia, PA, USA). A 10 mM stock solution was prepared for ONC201 with dimethylsulfoxide (DMSO) for all *in vitro* experiments. All stock solutions were stored in aliquots at −20 °C. For *in vivo* studies, ONC201 was freshly prepared with sterile saline at a concentration of 5.5 mg/mL before injection.

### Cell culture

Primary human glioma cell lines were established as previously described ^39^. A description of the cell lines can be found in **Table 1**. The GL261 murine glioma cell line was obtained from Charles River Laboratories, Inc., Frederick, MD. GL261 cells were cultured in log-growth phase in DMEM (Invitrogen, Carlsbad, CA) (supplemented with 10% fetal bovine serum, penicillin and streptomycin). Primary glioblastoma cells were grown under serum-free conditions in ultra-low adhesion plates in DMEM/F12, supplemented with B27, EGF, bFGF and heparin as described previously. All cells were grown in a humidified atmosphere at 37°C with 5% CO_2_. The unique identity of all patient-derived specimen was confirmed by DNA fingerprinting (Laragen, Culver City, CA). All lines were routinely tested for mycoplasma infection (MycoAlert, Lonza).

### Animals

6–8-week-old C57BL/6 mice, or NOD-*scid* IL2Rgamma^null^ (NSG) originally obtained from The Jackson Laboratories (Bar Harbor, ME) were re-derived, bred and maintained in a pathogen-free environment in the American Association of Laboratory Animal Care-accredited Animal Facilities of Department of Radiation Oncology, University of California, Los Angeles, in accordance to all local and national guidelines for the care of animals. Weight of the animals was recorded every day. 2×10^5^ GL261-Luc or 3×10^5^ HK-374-Luc cells were implanted into the right striatum of the brains of mice using a stereotactic frame (Kopf Instruments, Tujunga, CA) and a nano-injector pump (Stoelting, Wood Dale, IL). Injection coordinates were 0.5 mm anterior and 2.25 mm lateral to the bregma, at a depth of 3.0 mm from the surface of the brain. Tumors were grown for 3 days with successful grafting confirmed by bioluminescence imaging. Mice that developed neurological deficits requiring euthanasia were sacrificed.

### *In-vitro* sphere formation assay

For the assessment of self-renewal *in vitro*, primary GBM cells were irradiated with 0, 2, 4, 6 or 8 Gy and seeded under serum-free conditions into non-tissue-culture-treated 96-well plates in DMEM/F12 media, supplemented with 10 mL / 500 mL of B27 (Invitrogen), 0.145 U/mL recombinant insulin (Eli Lilly, Indiana), 0.68 U/mL heparin (Fresenius Kabi, Illinois), 20 ng/mL fibroblast growth factor 2 (bFGF, Sigma) and 20 ng/mL epidermal growth factor (EGF, Sigma). The number of spheres formed at each dose point was normalized against the non-irradiated control. The resulting data points were fitted using a linear-quadratic model.

### Secondary and tertiary glioblastoma spheres

Primary HK-308 and HK-374 spheres were grown in 10-cm petri dishes and treated with 2.5 μM ONC201 one hour before a single dose of 4 Gy radiation. The irradiated spheres were dissociated after 4-5 and plated at clonal densities to generate secondary spheres. Re-plated secondary spheres were treated with a single dose of ONC201 (2.5 μM) 24 hours later, dissociated, and plated at clonal densities after 4-5 days in suspension and cultured to form tertiary spheres.

### Clonogenic assay

GBM HK-374 cells were trypsinized and plated at a density of 100 cells per well in a 6-well plate for 0 and 2 Gy, 200 cells per well for 4 and 6 Gy, and 400 cells per well for 8 Gy. ONC201 was applied one hour before irradiation at a concentration of 500 nM, 1 μM, 2.5 μM and 5 μM. After 10 days, the colonies were fixed and stained with 0.1% crystal violet. Colonies containing at least 50 cells were counted in each group and presented as the percentage to the initial number of cells plated.

### Cell viability assay

GBM HK-374 cells were trypsinized and plated at a density of 1 x 10^5^ cells per well in a 6-well ultra-low adhesion plate. ONC201 was applied one hour before a single dose of irradiation (0 or 8 Gy) at a concentration of 500 nM, 1 μM, 2.5 μM and 5 μM. The spheres were collected and dissociated after 24, 48 and 72 hours. Live, trypan blue-excluding cells were counted using a hemocytometer and normalized to the cell counts of the 0 Gy DMSO-treated group.

### In vivo bioluminescent imaging

The mice were injected intraperitoneally with 100 μl of D-luciferin (15 mg/ml, Gold biotechnology). Five minutes later, animals were anesthetized (3% isoflurane gas in O_2_) and luminescence was recorded (IVIS Spectrum, Perkin Elmer, Waltham, MA). Images were analyzed with Living Image Software (Caliper Life Sciences).

### Drug treatment

After confirming tumor grafting via bioluminescent imaging, mice bearing GL261 tumors or HK-374 specimen were injected intraperitoneally (i.p.) on a weekly basis either with ONC201 (50 mg/kg) or saline until they reached the euthanasia endpoint. ONC201 was dissolved in sterile saline at a concentration of 5.5 mg/ml.

### Irradiation

Cells were irradiated at room temperature using an experimental X-ray irradiator (Gulmay Medical Inc. Atlanta, GA) at a dose rate of 5.519 Gy/min for the time required to apply a prescribed dose. The X-ray beam was operated at 300 kV and hardened using a 4mm Be, a 3mm Al, and a 1.5mm Cu filter and calibrated using NIST-traceable dosimetry. Corresponding controls were sham irradiated.

All mice were irradiated using an image-guided small animal irradiator (X-RAD SmART, Precision X-Ray, North Branford, CT) with an integrated cone beam CT (60 kVp, 1 mA) and a bioluminescence-imaging unit. The X-ray beam was operated at 225 KV and calibrated with a micro-ionization chamber using NIST-traceable dosimetry. Cone beam CT images were acquired with a 2 mm Al filter. For delivery of the radiation treatment the beam was hardened using a 0.3 mm Cu filter. During the entire procedure the interior of the irradiator cabinet was maintained at 35° C to prevent hypothermia of the animals. Anesthesia of the animals was initiated in an induction chamber. Once deeply anesthetized, the animals were then immobilized using custom 3D-printed mouse holder (MakerBot Replicator+, PLA filament) with ear pins and teeth bar, which slides onto the irradiator couch. Cone beam CT images were acquired for each individual animal. Individual treatment plans were calculated for each animal using the SmART-Plan treatment planning software (Precision X-Ray). Radiation treatment was applied using a square 1 cm x 1 cm collimator from a lateral field. For the assessment of the effect of ONC201 in combination with irradiation *in vivo*, mice were treated with ONC201 (50 mg/kg, i.p.) or saline one hour prior to irradiation. Animals received a single dose of 10 Gy on day 3 after tumor implantation.

### Immunohistochemistry

Brains were explanted, fixed in formalin for twenty-four hours and embedded in paraffin. 4 μm sections were stained with hematoxylin and eosin (H&E) using standard protocols. Additional sections were baked for 30 minutes in an oven at 65 °C, dewaxed in two successive Xylene baths for 5 minutes each and then hydrated for 5 minutes each using an alcohol gradient (ethanol 100%, 90%, 70%, 50%, 25%). Antigen retrieval was performed using Heat Induced Epitope Retrieval in a citrate buffer (10 mM sodium citrate, 0.05% tween20, pH 6) with heating to 95 °C in a steamer for 20 minutes. After cooling down, the slides were blocked with 10% goat serum plus 1% BSA at room temperature for 30 minutes and then incubated with the primary antibody against Ki67 (Cell signaling, Cat #12202S, 1:800) or c-Myc (Cell signaling, Cat #5605S, 1:800) overnight at 4 °C. The next day, the slides were rinsed with PBS and then incubated with ready-to-use IHC detection reagent (Cell signaling, Danvers, MA; 10 μl) at room temperature for one hour, rinsed, and then incubated with DAB (Cell Signaling) for 3-5 minutes. Tissues were counterstained with Harris modified Hematoxylin (Fisher scientific, Waltham, MA) for 30 seconds, dehydrated via an alcohol gradient (ethanol 25%, 50%, 70%, 90%, 100%) and soaked twice into Xylene. A drop of Premount mounting media (Fisher Scientific) was added on the top of each section before covering up with a coverslip.

### Quantitative Reverse Transcription-PCR

Total RNA was isolated using TRIZOL Reagent (Invitrogen). cDNA synthesis was carried out using the SuperScript Reverse Transcription IV (Invitrogen). Quantitative PCR was performed in the QuantStudio™ 3 Real-Time PCR System (Applied Biosystems, Carlsbad, CA, USA) using the PowerUp™ SYBR™ Green Master Mix (Applied Biosystems). *C*_t_ for each gene was determined after normalization to IPO8, TBP, PPIA and ΔΔ*C*_t_ was calculated relative to the designated reference sample. Gene expression values were then set equal to 2^−ΔΔCt^ as described by the manufacturer of the kit (Applied Biosystems). All PCR primers were synthesized by Invitrogen with PPIA, TBP and IPO8 as housekeeping genes (for primer sequences see **Supplementary Table1**).

### RNA-Sequencing

Primary HK-374 cells were treated with 2.5 μM ONC201 one hour before a single dose of radiation (4 Gy), RNA was extracted at 48 hours using Trizol followed with RNeasy kit (Qiagen) isolation. The next-generation sequencing was performed by Novogene (Chula Vista, CA). Quality and integrity of total RNA was quantified on the Agilent Technologies 2100 Bioanalyzer (Agilent Technologies; Waldbronn, Germany). The sequencing and analysis process was performed as previously reported^40^.

Briefly, differential expression analysis between irradiated and control samples (three biological replicates per condition) was performed using the DESeq2 R package (2_1.6.3). The resulting *p*-values were adjusted using the Benjamin and Hochberg’s approach for controlling the False Discovery Rate (FDR). Genes with an adjusted *p*-value of <0.05 found by DESeq2 were assigned as differentially expressed. To identify the correlations between differentially expressed genes (DEGs), we generated heatmaps using the hierarchical clustering distance method with the function of heatmap, SOM (Self-organization mapping) and k-means using the silhouette coefficient to adapt the optimal classification with default parameters in R software.

Gene Ontology (GO) enrichment analysis of differentially expressed genes was generated using the **GO**rilla (**G**ene **O**ntology en**R**Ichment ana**L**ysis and vizua**L**iz**A**tion) tool. GO terms with corrected *p*-values less than 0.05 were considered significantly enriched by differentially expressed genes.

Gene set enrichment analysis (GSEA) using all genes ranked by their differential expression as input was performed for *Hallmark* and *Curated* gene sets (http://software.broadinstitute.org/gsea/index.jsp; accessed 05/2020). GSEA with gene sets related to quiescent glioblastoma signatures were manually searched by the gene sets list and ECM-related genes were sorted out for analysis^35^.

### TCGA data mining and analysis

The TCGA Provisional dataset (captured June 1, 2020) was accessed via cBioPortal^41,42^. Kaplan-Meier estimates were calculated for patients with upregulation (z-score >1.5) in one or more genes enriched in qGBM cells (FN1, CAV1, COL12A1, EFNB2, AKAP12, TNC, CDH13, SPARC, COL6A2, ITGA3, COL4A1, AHNAK, CD44, CTSB, LAMA4, CD59, SVIL, VCL, LAMC1, COL15A1, ACTN1, SYNE2) and compared to patients with no alterations in the expression of those genes.

### Statistics

Unless stated otherwise all data shown are represented as mean ± standard error mean (SEM) for at least 3 biologically independent experiments. A *p*-value of ≤0.05 in an unpaired two-sided Student’s t-test or Two-way ANOVA test indicated a statistically significant difference. Kaplan-Meier estimates were calculated using the GraphPad Prism Software package. For Kaplan-Meier estimates a *p*-value ≤0.05 in a log-rank (Mantel-Cox) test indicated a statistically significant difference.

### Data sharing

All data and methods are included in the manuscript. Patient-derived cell lines will be made available upon reasonable request. The RNA-Seq data are available via GEO accession number GSE153982.

## Results

### ONC201 decreases cell viability, clonogenicity and self-renewal in GBM

ONC201 was originally discovered by screening drug libraries for compounds that induce TRAIL to initiate death receptor-mediated apoptosis. To study the effects of ONC201 on GBM, exponentially growing HK-374 cells were treated with ONC201 at 0.5, 1, 2.5 or 5 μM, concentrations known to be reached in the brain lesions of patients with GBM^21^, and irradiated with 0 or 8 Gy. At 24, 48 and 72 hours after the start of the treatment, we observed a significant, dose-dependent reduction in the number of viable cells, thus suggesting cytostatic or cytotoxic activity of ONC201 on the bulk population of HK-374 GBM cells (**Fig. 1A**).

**Figure 1.**
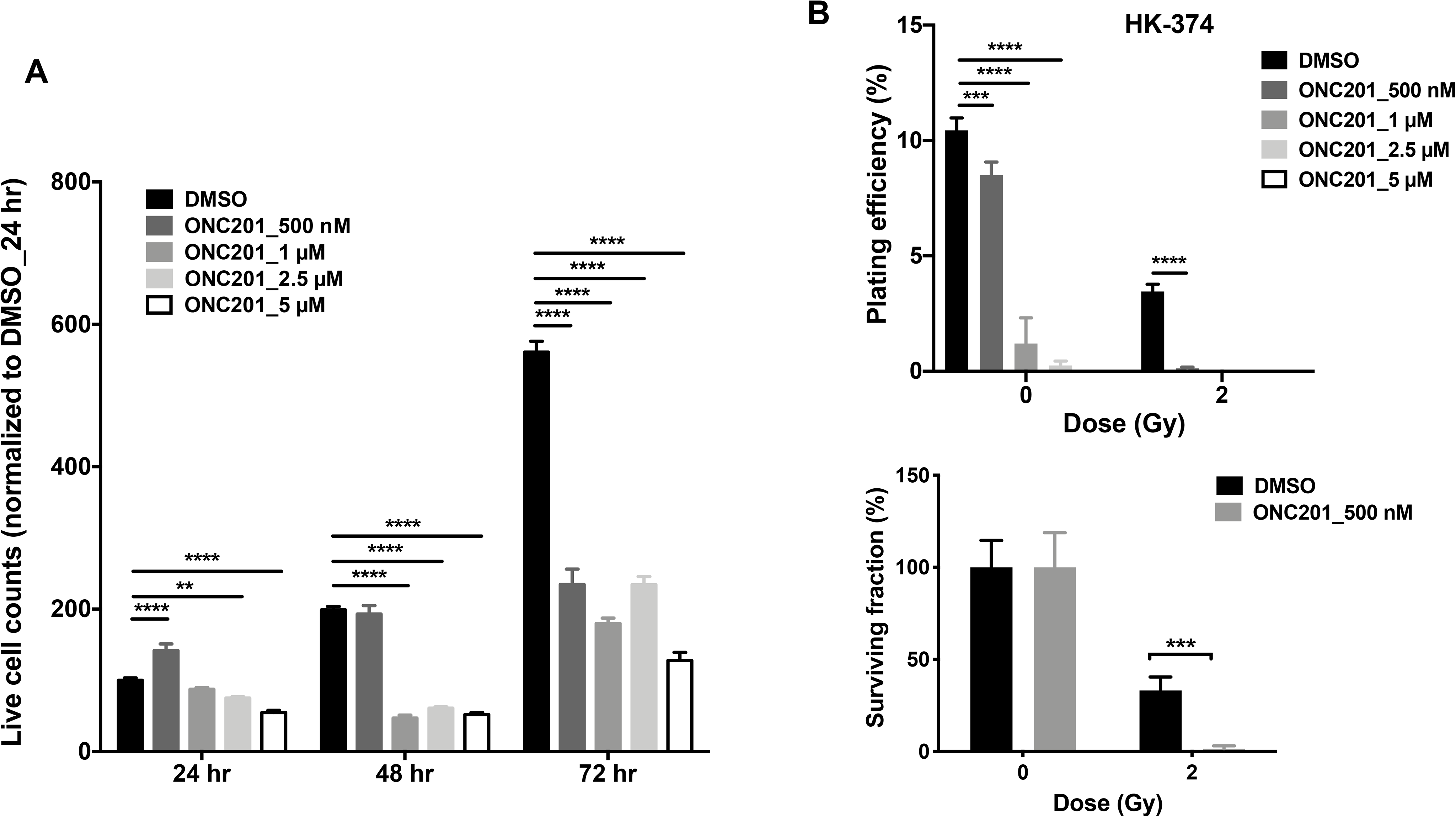
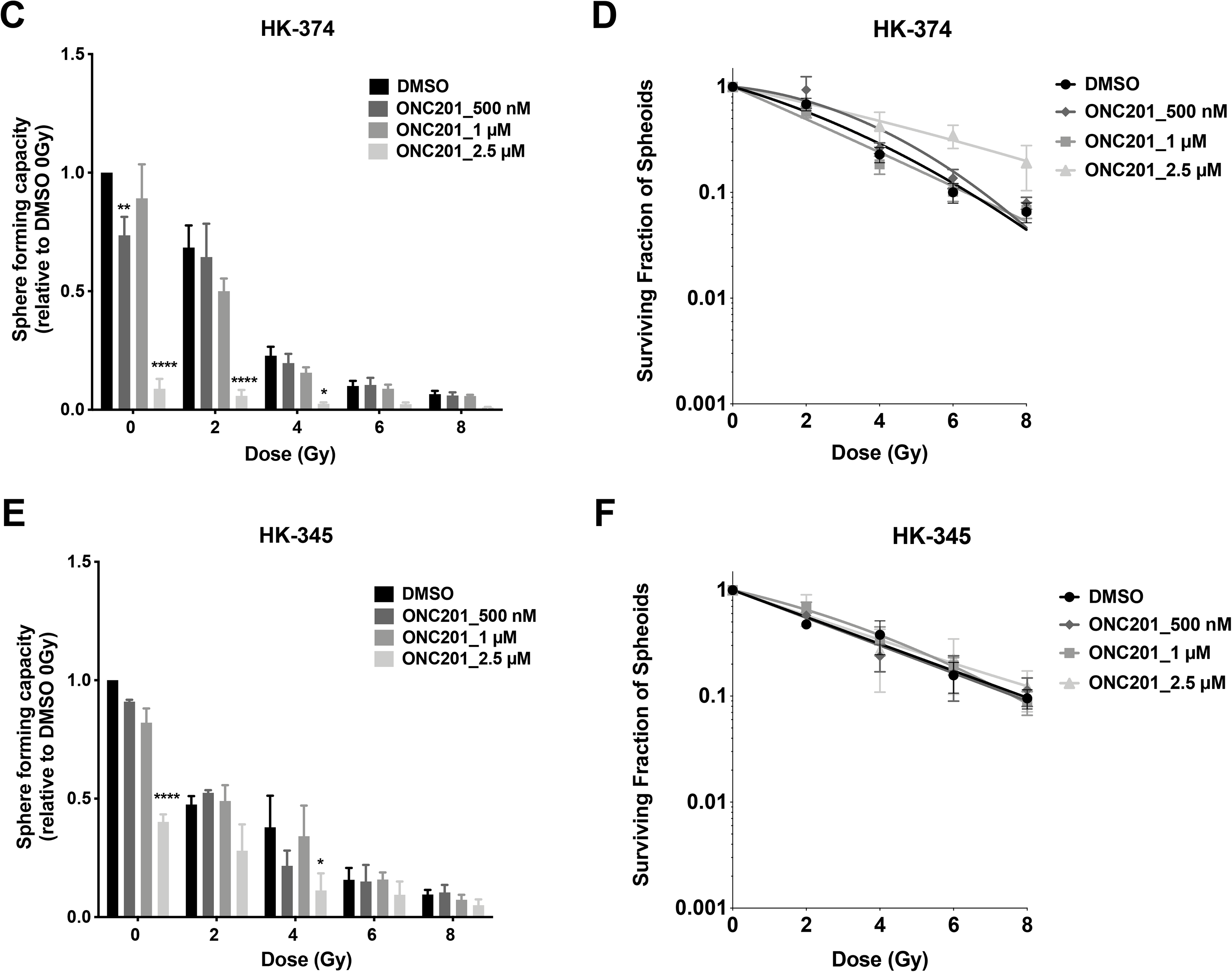
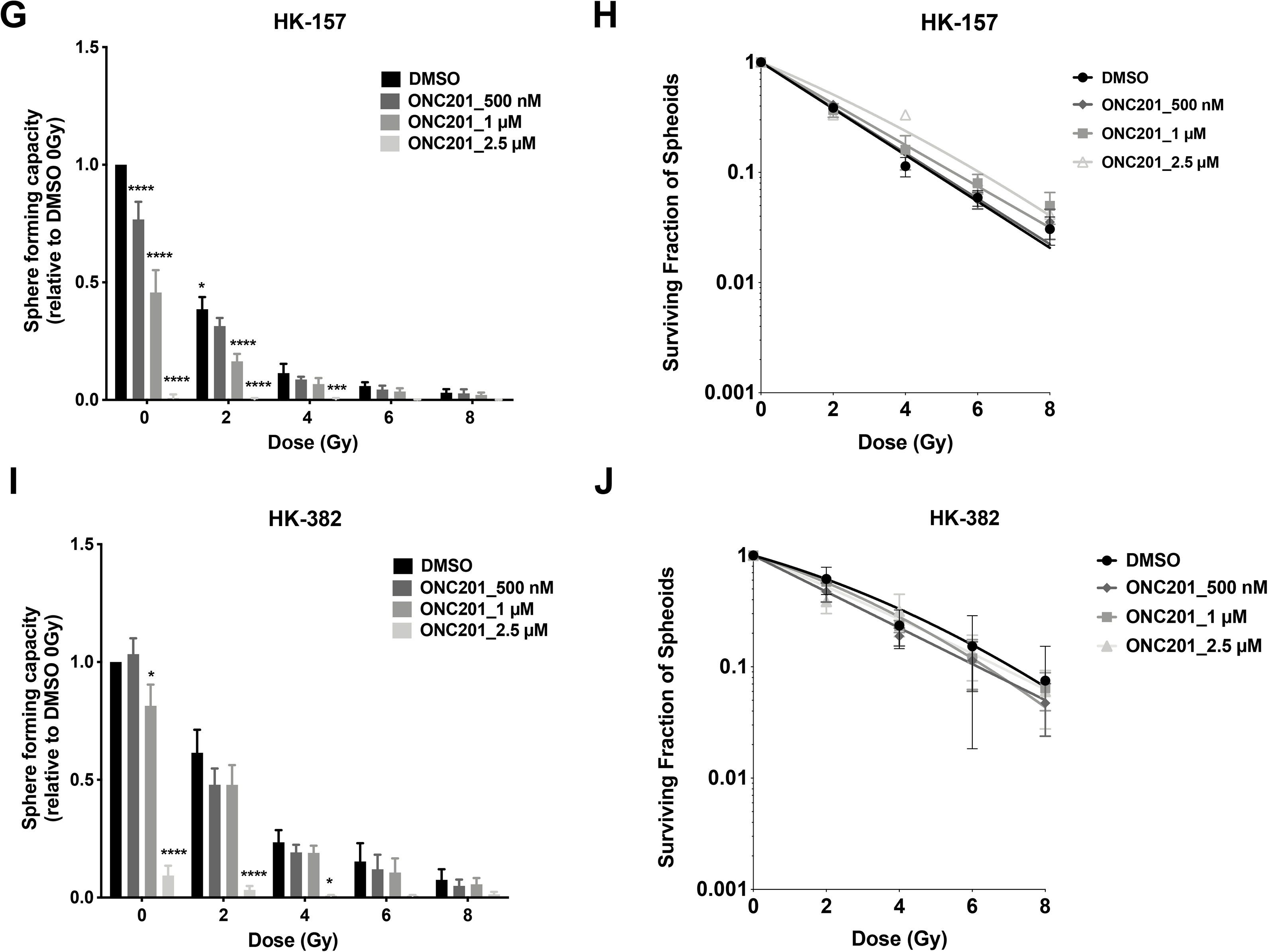
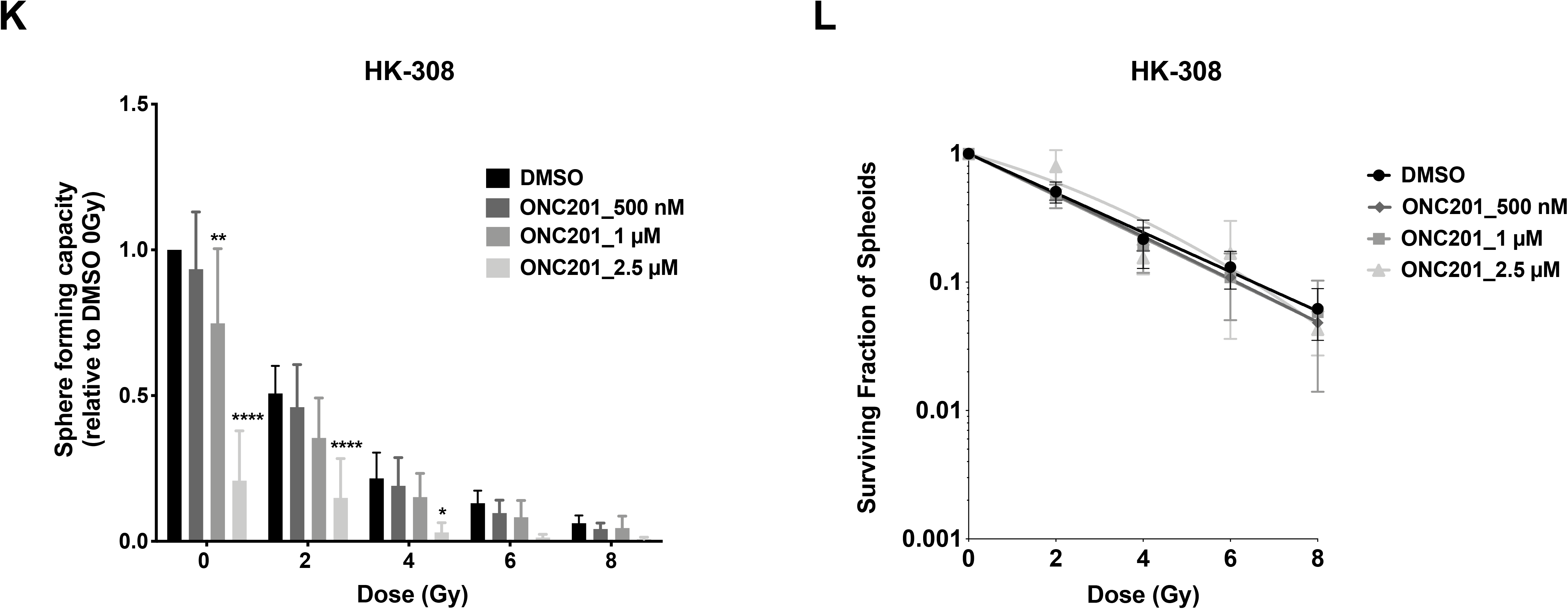
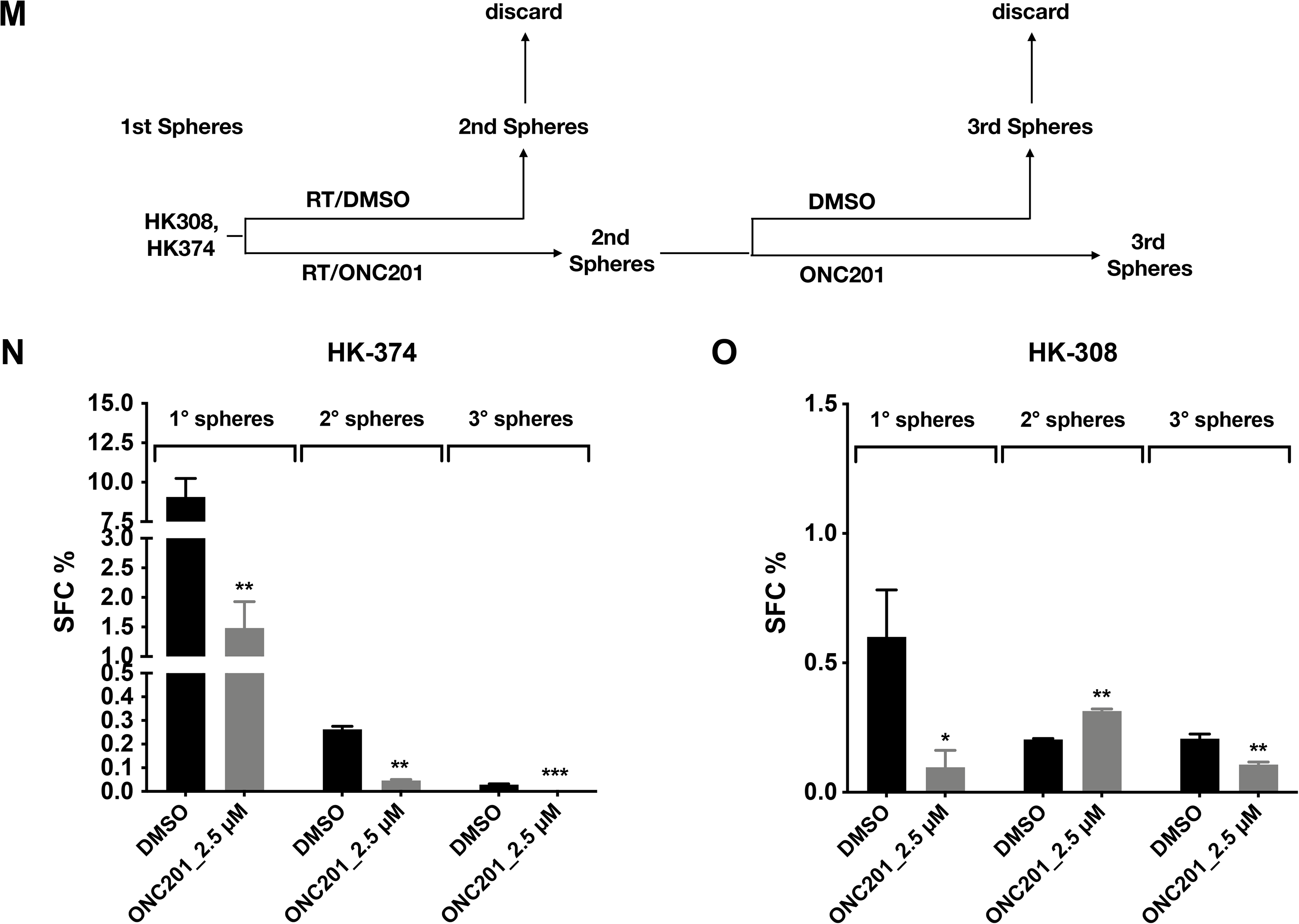
Effects of ONC201 on GBM cells *in vitro*. (**A**) Live cell counts of HK-374 cells treated with ONC201 (500 nM, 1 μM, 2.5 μM and 5 μM) or solvent control (DMSO) for 24, 48 or 72 hours. (**B**) Clonogenic assay of patient-derived primary HK-374 GBM cells treated with ONC201 (500 nM, 1 μM, 2.5 μM and 5 μM) or solvent control (DMSO) in combination with a single dose of radiation (0 or 2 Gy). The colony number was counted and presented as the percentage relative to the initial number of cells plated. (**C-L**) Patient-derived HK-374, HK-345, HK-157, HK-382 and HK-308 GBM cells were used to perform sphere-forming assays with sham-irradiated or irradiated cells in the presence or absence of ONC201 (500 nM, 1 μM, 2.5 μM). The spheres were cultured in suspension for 7-10 days. The number of spheres formed under each condition were counted and presented as percentage spheres formed and normalized against the sham-irradiated control. (**M**) Schematic flow chart of secondary and tertiary spheres from patient-derived HK-308 and HK-374 GBM cells. (**N/O**) Sphere-forming capacity (SFC%) was evaluated using HK-374 and HK-308 derived secondary and tertiary spheres in the presence or absence of 2.5 μM ONC201 using a limiting dilution assay. All experiments have been performed with at least 3 biological independent repeats. *p*-values were calculated using two-way ANOVA for A-L; multiple Student’s t-tests for N and O. * *p*-value<0.05, ** *p*-value<0.01, *** *p*-value<0.001 and **** *p*-value<0.0001.

A radiobiological gold-standard for assessing the radiosensitizing properties of drugs is the clonogenic survival assay, which measures the formation of adherent colonies of more than 50 cells from an individual clonogenic cell. Treatment of HK-374 cells with ONC201 led to a significant, dose-dependent reduction in the plating efficiency of HK-374 cells. Normalizing to the plating efficiency at 0 Gy, 0.5 μM concentrations of ONC201 caused a drop in the surviving fraction of the cells from 33.1 ± 3% to 1.5 ± 0.7% (*p*=1.2×10^−6^, Student’s t-test). Higher concentrations of ONC201 in combination with 2 Gy prevented colony formation completely (**Fig. 1B**).

Gliomaspheres are known to enrich for glioma-initiating cells and gliomasphere formation from a single cell is a functional measure of self-renewal capacity^43,44^. To assess the effect of ONC201 on self-renewal capacity, we first performed an *in vitro* limiting dilution assay using the patient-derived GBM lines HK-382, HK-374, HK-157, HK-345 and HK-308. Cells were treated with a single dose of radiation (0, 2, 4, 6 or 8 Gy) in the presence or absence of ONC201 (0, 0.5, 1 or 2.5 μM). Treatment with ONC201 led to a dose-dependent reduction in sphere-formation (**Fig. 1C/E/G/I/K**). Curve fitting of the sphere-forming data using a linear-quadratic model indicated that ONC201 did not act as a classical radiosensitizer for glioma-initiating cells (**Fig. 1D/F/H/J/L**).

Radiation therapy prolongs the survival of patients with GBM, but the tumors almost always recur. In order to test if ONC201 inhibits self-renewal of GBM cells surviving combined treatment of a sublethal dose of radiation and ONC201 we next performed secondary and tertiary sphere formation assays in the absence and presence of ONC201 using cells that survived a combination of 4 Gy and ONC201 (2.5 μM) (**Fig. 1M**). Irradiated, ONC201-treated cells displayed reduced primary, secondary and tertiary sphere formation in both, the HK-374 line (established from a primary GBM) (**Fig. 1N**) and the HK-308 line (established from a recurrent GBM) (**Fig. 1O**) when compared to primary spheres formed from cells treated with 4 Gy only.

When surviving HK-374 cells from primary spheres from the combined treatment with radiation and ONC201 were further treated with ONC201, the line showed exhaustion of sphere-forming cells in secondary and tertiary spheres (**Fig. 1N**). Primary sphere-formation in the HK-308 line was more than an order of magnitude lower than in the HK-374 line. However, sphere-formation after additional rounds of ONC201 treatment did not indicate further loss of sphere-forming capacity in secondary or tertiary spheres (**Fig. 1O**).

### Gene signature profiles of GBM cells in response to radiation and ONC201

In order to better understand the diverse effects of ONC201 on different populations of GBM cells, including the reduction in viability of bulk cell populations, the loss in clonogenicity and radiosensitization of clonogenic cells as well as the diminished self-renewal of glioma-initiating cells, we performed RNA-Sequencing (RNA-Seq) on HK-374 cells, 48 hours after irradiation and/or ONC201 treatment. Principal component analysis (PCA) showed that biologically-independent replicates of each experimental group clustered closely, indicating reproducibility of the results (**Fig. 2A**). Hierarchical clustering of differentially expressed genes revealed a distinct gene expression profile after combined treatment with radiation and ONC201 (**Fig. 2B**). Radiation caused differential upregulation of 1711 genes and down-regulation of 1680 genes when compared to DMSO control, while the combination of ONC201 and radiation led to 3048 differentially up-regulated and 3156 down-regulated genes when compared to radiation only. We also compared combination-treated samples and ONC201-treated samples to DMSO-treated controls, and found 3190 differentially up-regulated and 3158 down-regulated genes in combination-treated, and 2989 differentially up-regulated and 3223 down-regulated genes in ONC201-treated cells (**Fig. 2C**). The top 10 up- and down-regulated genes with their log2 fold changes for all four comparisons are shown in **Fig. 2D**.

**Figure 2.**
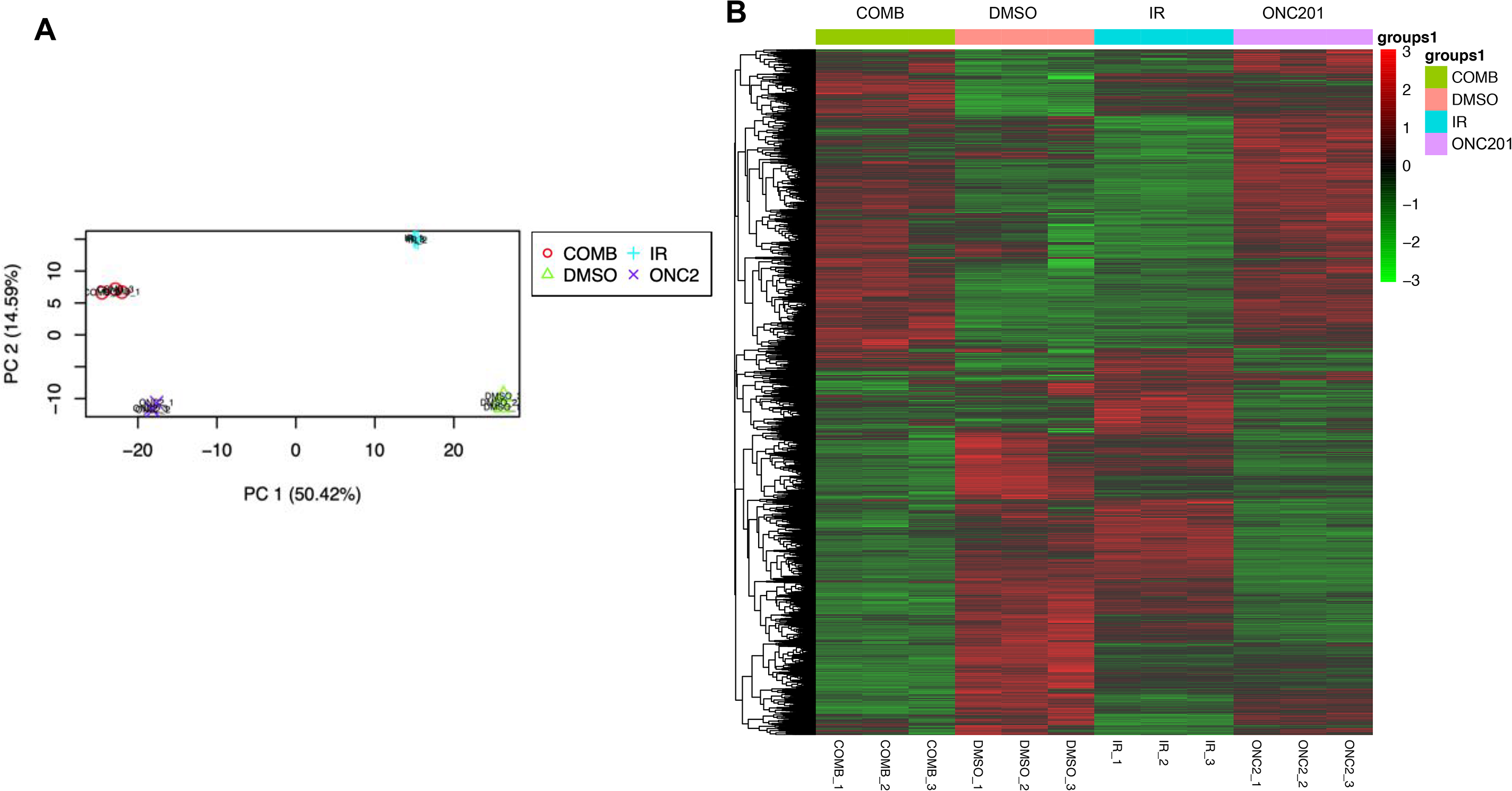
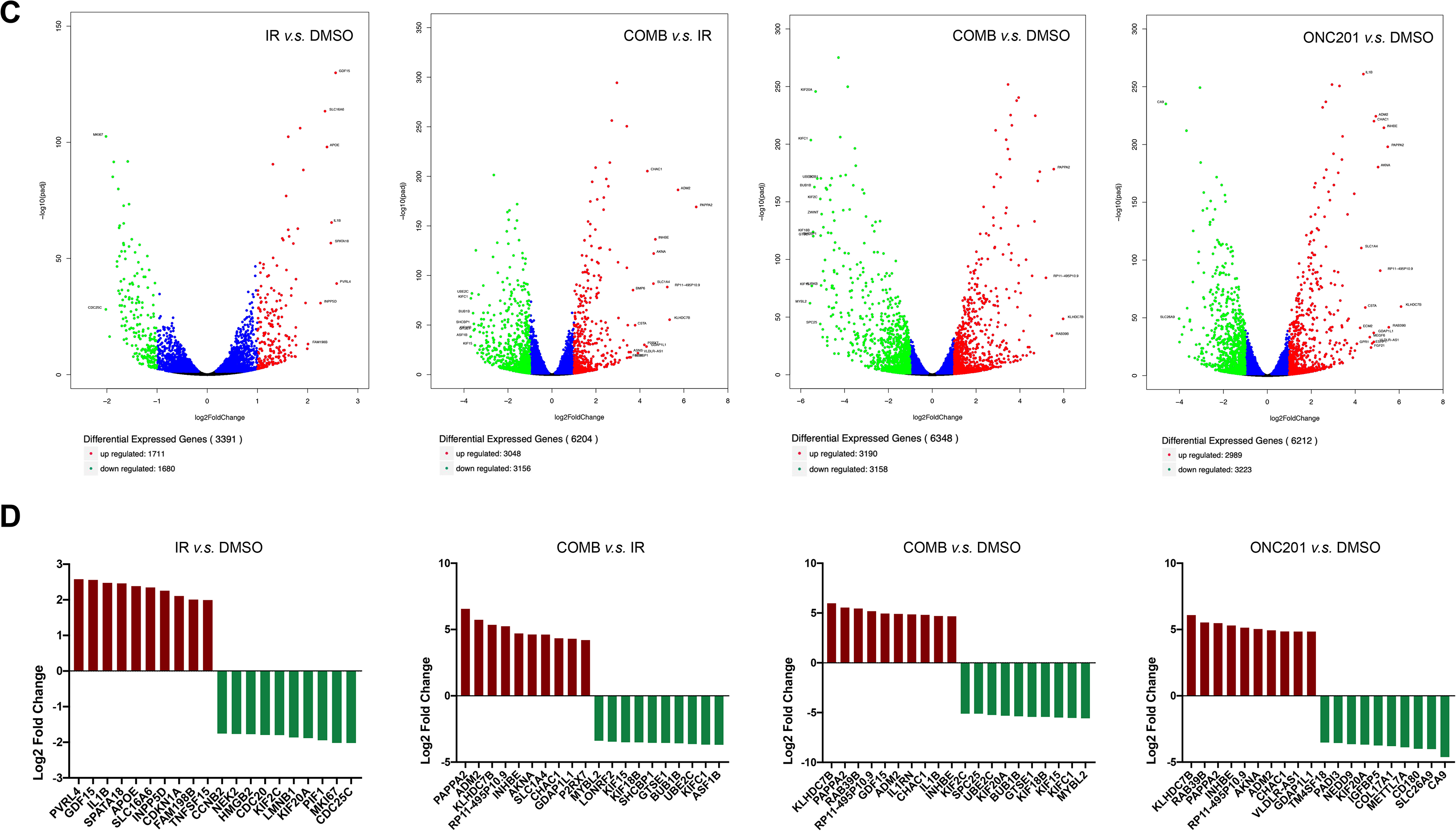
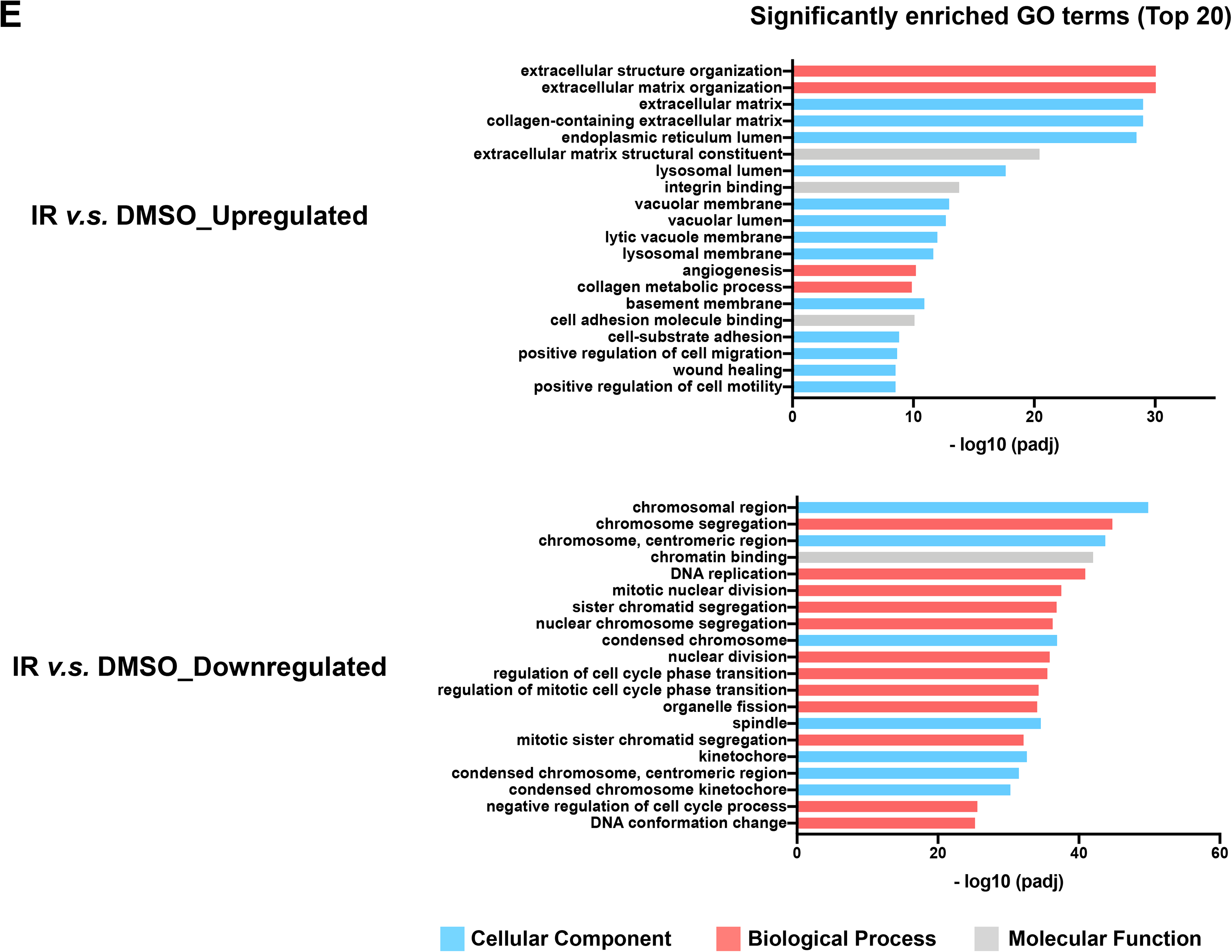
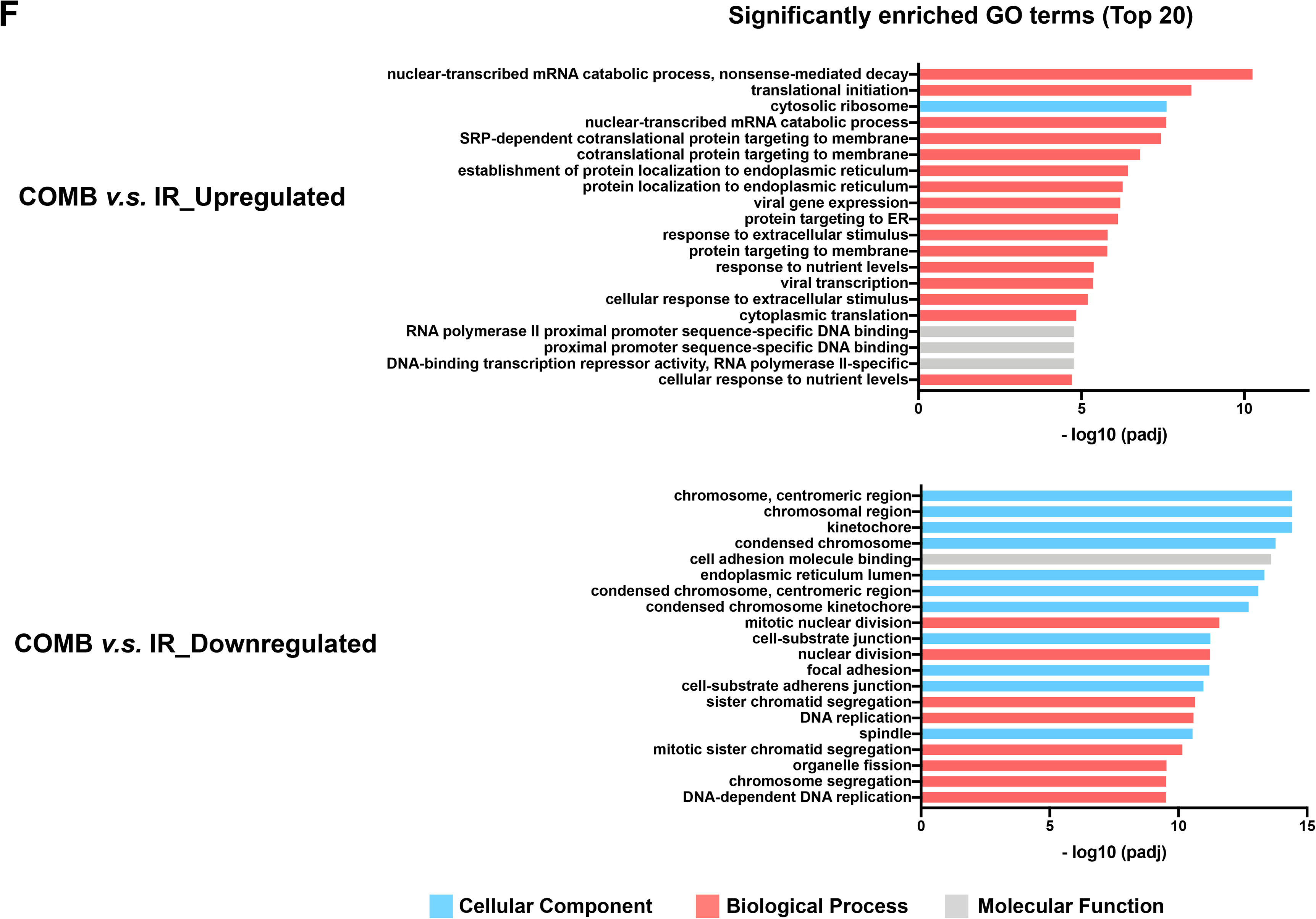
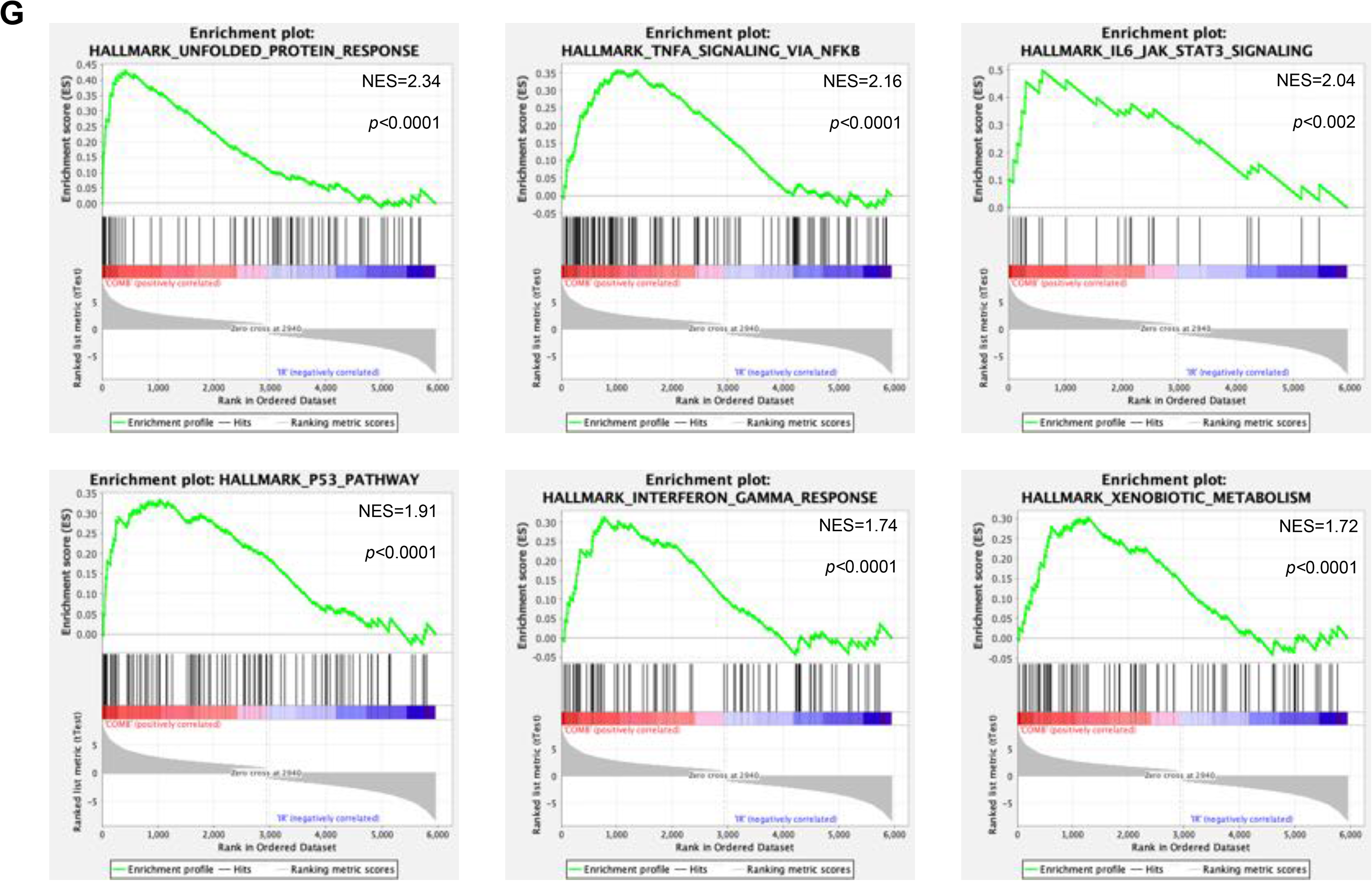
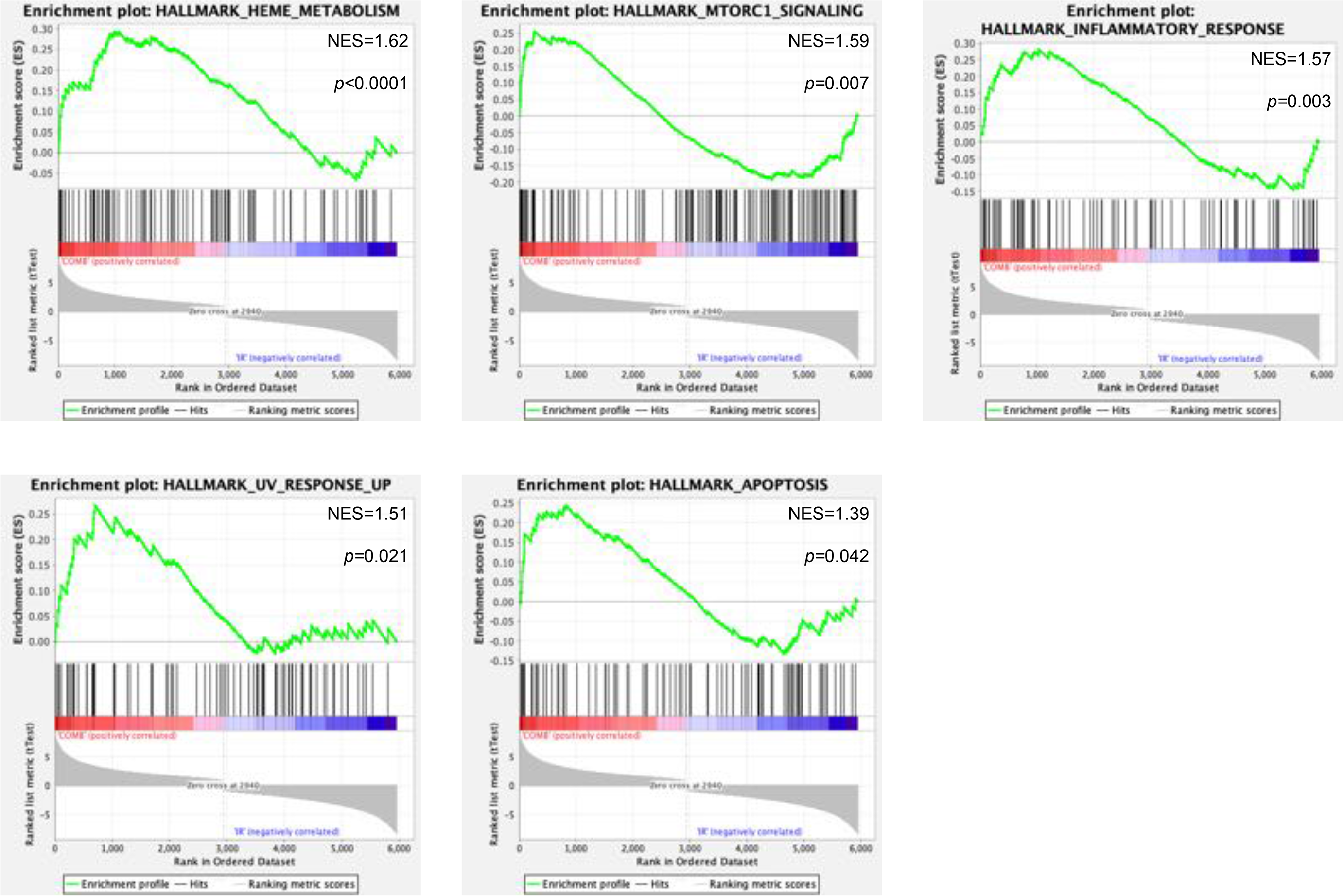
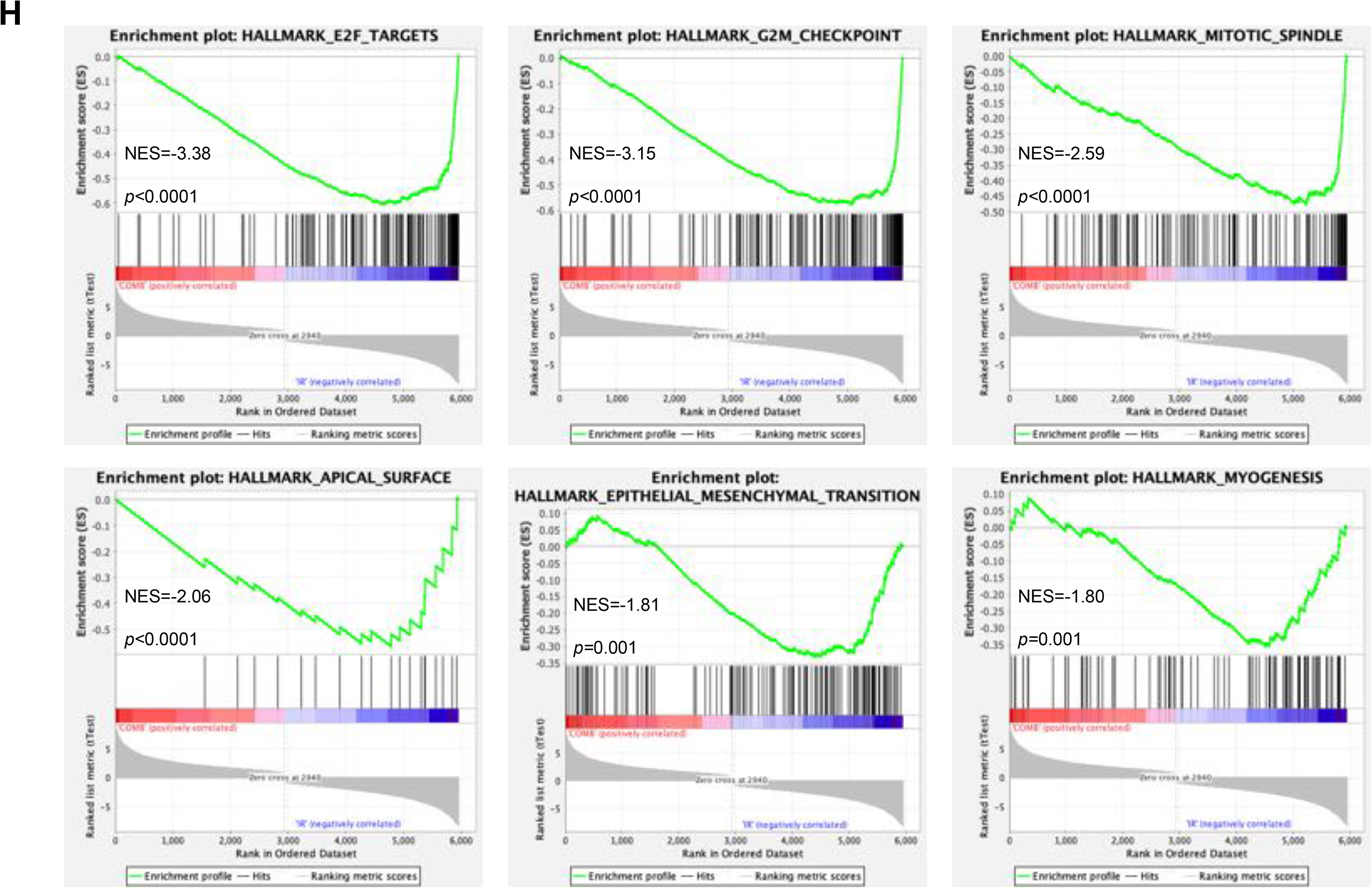
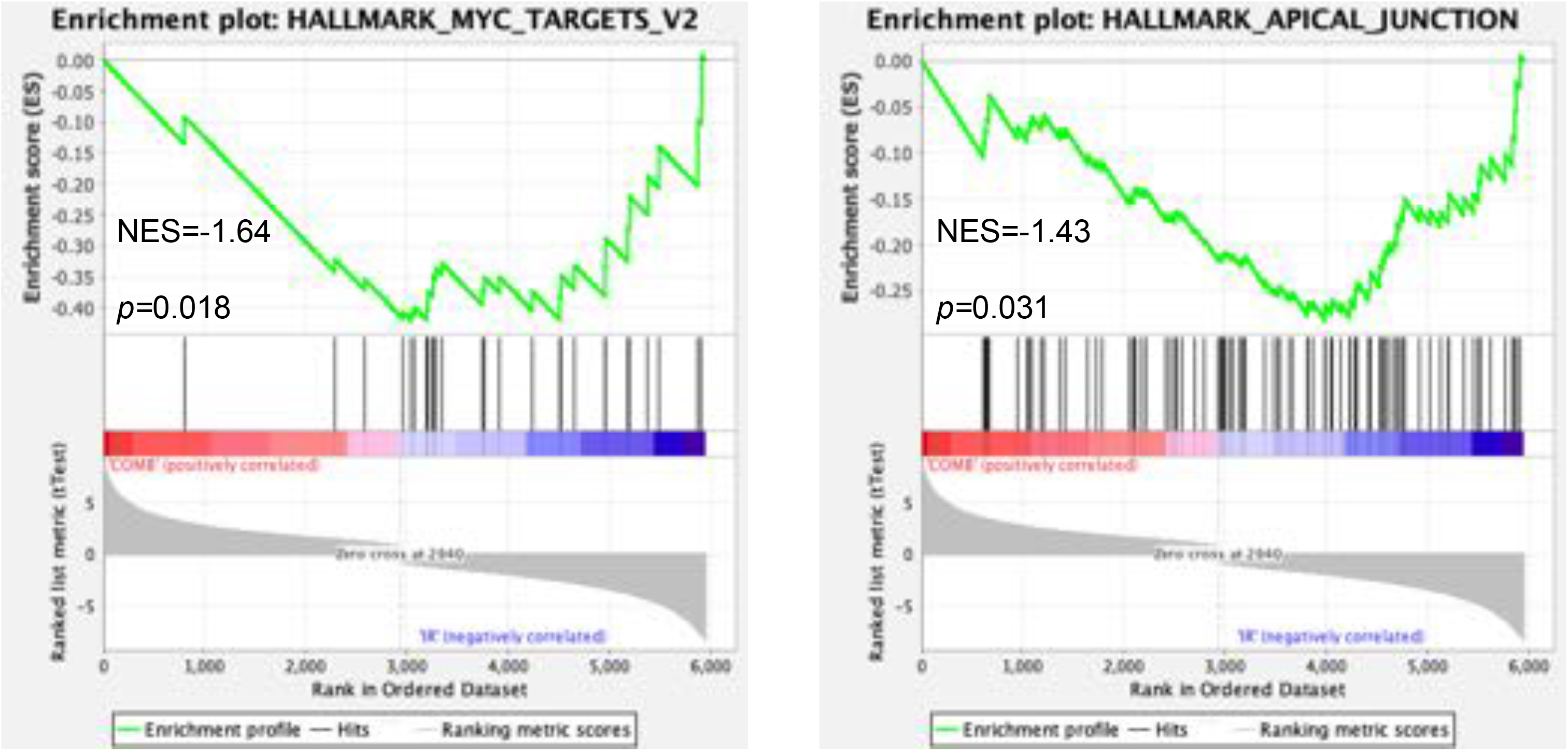
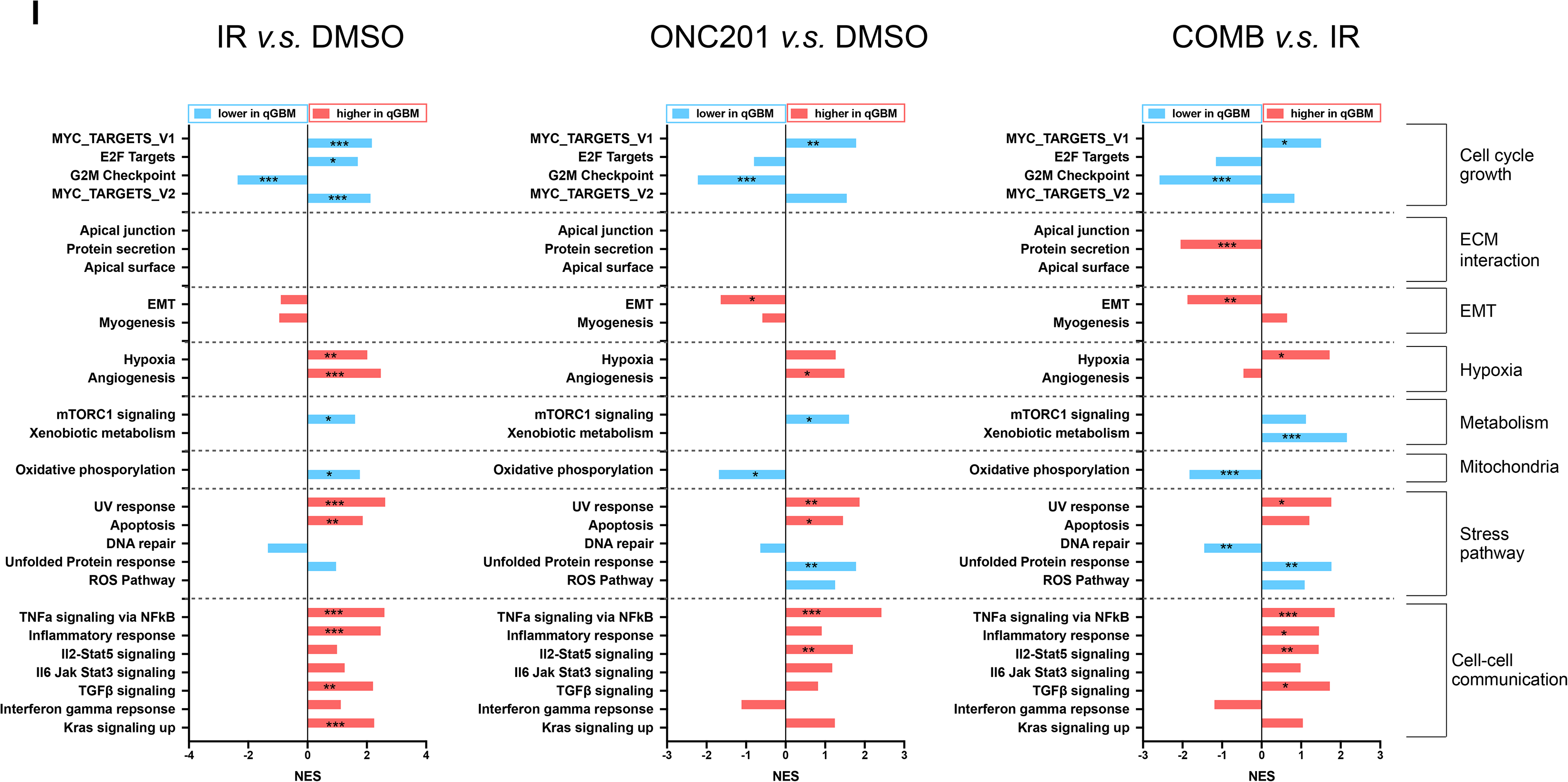
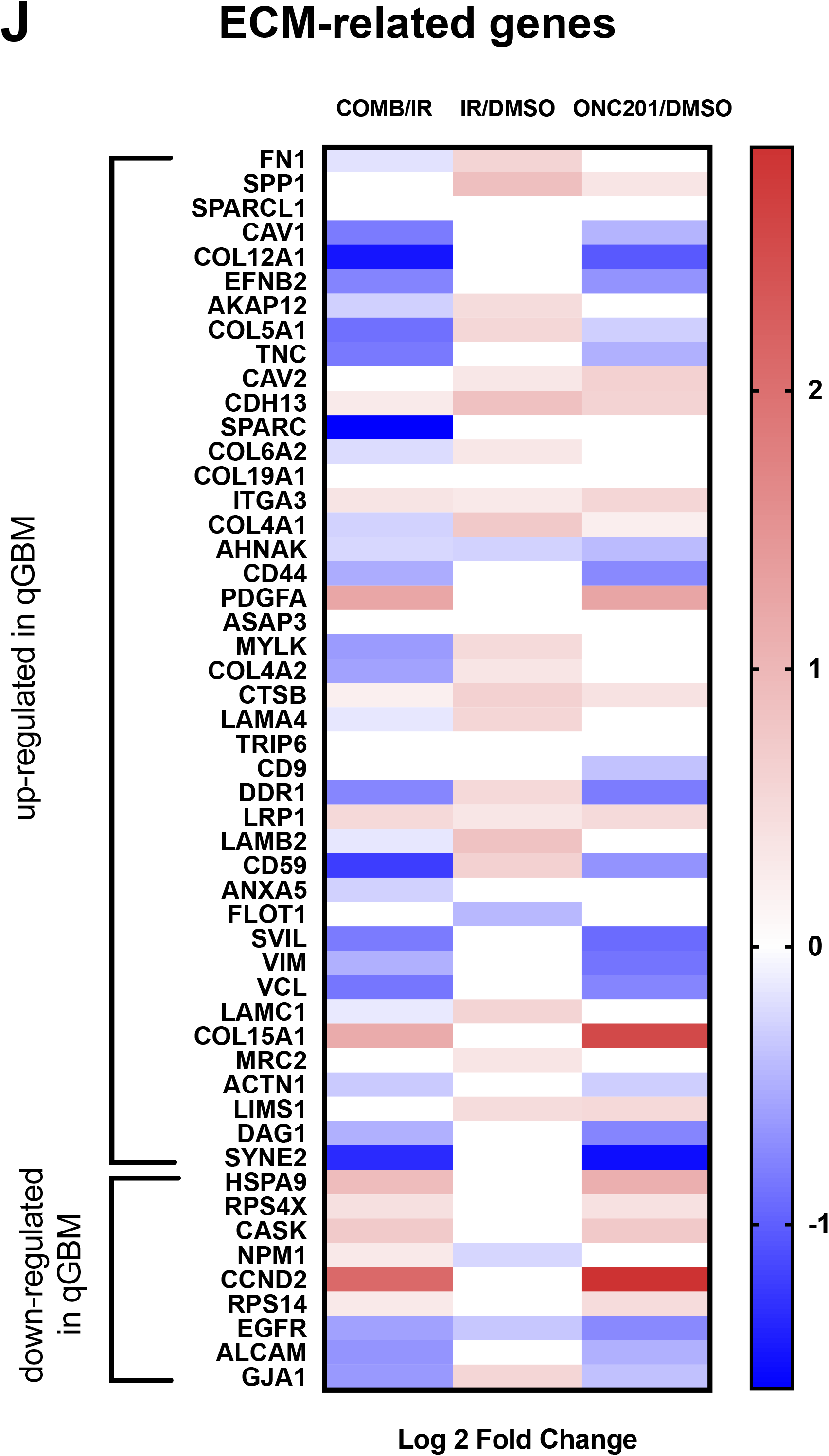
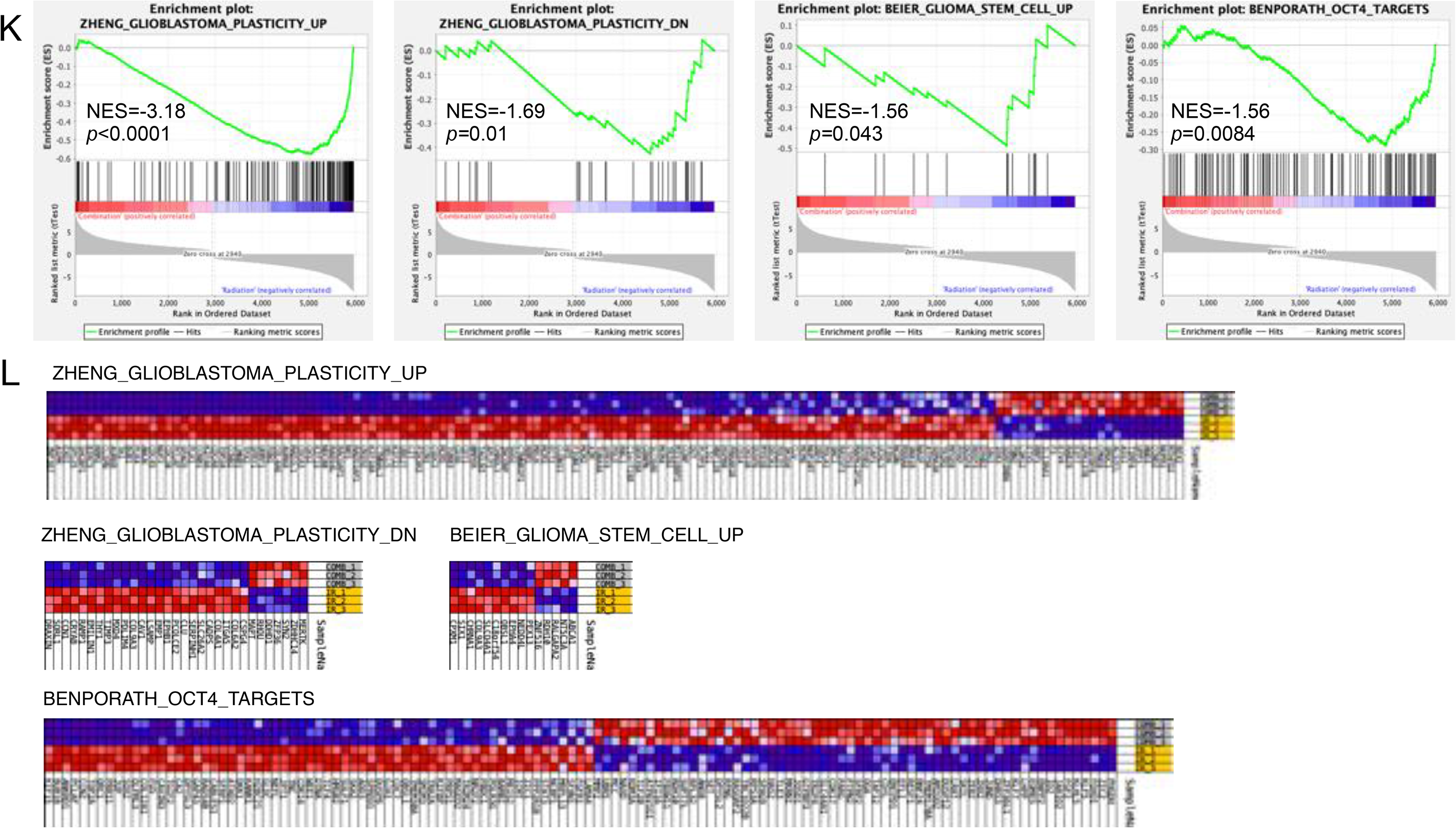
RNA-Seq analysis of glioma cells treated with ONC201 and radiation. (**A**) Principal Component Analysis (PCA) plot displaying all 12 samples along PC1 and PC2, which describe 50.42% and 14.59% of the inter- and intra-group variability respectively, within the expression dataset. PCA was applied to normalized (reads per kilobase of transcript per 1 million mapped reads) and log-transformed count data. (**B**) Heatmap of differentially expressed genes in HK-374 cells, 48 hours after treatment with radiation (4 Gy), ONC201, COMB (ONC201 in combination with radiation) or DMSO control. (**C**) Volcano diagrams of differentially expressed genes in HK-374 cells, 48 hours after treatment with radiation, ONC201 or COMB compared to DMSO-treated control cells. (**D**) Top 10 up- and down-regulated genes with their Log2 fold change in the comparisons, e.g., IR *v.s*. DMSO, COMB *v.s*. IR, COMB *v.s*. DMSO, ONC201 *v.s*. DMSO. (**E**) The top 20 significantly enriched signaling pathways in the Gene Ontology (GO) database between IR and DMSO. Up-regulated in IR *v.s*. DMSO (top) and down-regulated in IR *v.s*. DMSO (bottom). (**F**) The top 20 significantly enriched signaling pathways in the Gene Ontology (GO) database between COMB and IR. Up-regulated in COMB *v.s*. IR (top) and down-regulated in COMB *v.s*. IR (bottom). (**G/H**) GSEA results of the gene sets that are positively or negatively enriched in the combination of ONC201 and radiation as compared to radiation alone. NES, normalized enrichment score. (**I**) GSEA of quiescent GBM (qGBM) gene signatures in the comparisons IR *v.s*. DMSO, COMB *v.s*. IR, and ONC201 *v.s*. DMSO, displayed with normalized enrichment scores. Red indicates the gene sets highly expressed in qGBM, while the blue represents gene sets with low expression in qGBM. The nominal *p*-value was labeled with *<0.05; **<0.01; ***<0.001. (**J**) Heatmap of ECM-related gene expressions in the comparisons, e.g., IR *v.s*. DMSO, COMB *v.s*. IR, ONC201 *v.s*. DMSO, stratified by whether these genes are up- and down-regulated in qGBM. (**K/L**) GSEA results of the gene sets of glioblastoma plasticity, glioma stem cells and targets of the developmental transcription factor Oct4 in the combined treatment of ONC201 and radiation (COMB) compared to radiation alone (IR). Enrichment plots and corresponding heatmaps displayed.

The top up- and down-regulated genes in cells after combined treatment with radiation and ONC201 compared to irradiated cells were validated using qRT-PCR (**Supplementary Fig. 1**).

Next, we performed a GO enrichment analysis of DEGs in irradiated versus control samples. Differentially up-regulated genes overlapped with gene sets containing extracellular matrix (ECM) components (**Fig. 2E, Top**), while down-regulated genes overlapped with gene sets involved in DNA replication, chromatin binding, and mitotic nuclear division (**Fig. 2E, Bottom**). DEGs in combination-treated versus irradiated cells, overlapped with gene sets involved in catabolism, cellular response to extracellular stimuli, and DNA-binding transcription repressor activity (**Fig. 2F, Top**), while down-regulated DEGs overlapped with gene sets involved in cell adhesion, mitotic nuclear division, and DNA replication (**Fig. 2F, Bottom**).

Thereafter, we assessed global gene expression changes in combination-treated cells relative to irradiated cells using GSEA. DEGs in cells treated with radiation and ONC201 revealed an overlap with the Hallmark gene sets of apoptosis and the p53 pathway gene sets (**Fig. 2G**), consistent with the pro-apoptotic properties of ONC201 and a p53 response to radiation. Furthermore, DEGs up-regulated in response to a combination of radiation and ONC201 overlapped with gene sets of the unfolded protein response, TNFα and IL-6/Stat3 and mTORC1 signaling, IFNγ response, xenobiotic and heme metabolism, genes up-regulated in response to UV and an inflammatory response (**Fig. 2G**). Down-regulated DEGs in cells treated with radiation and ONC201 overlapped with the gene sets of E2F target, G2M checkpoint, mitotic spindle, apical surface, epithelial mesenchymal transition (EMT) and myogenesis (**Fig. 2H**). Heat maps of genes contributing to the leading edge of each gene set are shown in **Supplementary Fig. 2**.

GBM are known for their intratumoral heterogeneity. A recent study reported the presence of a relatively quiescent cell population in GBM (qGBM) that exhibited self-renewal capacity comparable to its proliferative counterparts (pGBM) but showed higher therapeutic resistance and distinct gene expression signatures^35^. We had previously confirmed the presence of quiescent and proliferating cells with self-renewal capacity in GBM but also reported recruitment of qGBM cells into the cell cycle in response to radiation^40^. Analyzing our transcriptome data, we observed pattern changes in gene sets associated with cell cycle growth, EMT, metabolism, mitochondria, and stress pathway in response to radiation (**Fig. 2I, left**) consistent with the recruitment of quiescent GBM into the cell cycle. Treatment of the cells with ONC201 led to a down-regulation of genes involved in the G2/M checkpoint, EMT-related genes and genes involved in oxidative phosphorylation, consistent with ClpP agonism, while up-regulating angiogenesis, mTORC1 signaling, UV response, apoptosis, unfolded protein response, TNFα and IL2/Stat5 signaling (**Fig. 2I, middle**). Compared to radiation alone, the combination of radiation and ONC201 reduced the expression of genes involved in G2/M checkpoint control, protein secretion, EMT, oxidative phosphorylation and DNA repair, while increasing the expression of Myc target genes, hypoxia-related genes, genes involved in xenobiotic metabolism, UV response, TNFα, IL2/Stat5, TGFβ and genes of a general inflammatory response (**Fig. 2I, right**). A subpopulation of GBM cells is thought to acquire and maintain quiescence through ECM organization and interaction with niche factors^35^.

When compared to irradiated samples, the combined treatment of radiation and ONC201 significantly down-regulated genes involved in ECM interaction, while irradiation or treatment with ONC201 alone did not significantly affect the expression of these genes (**Fig. 2J**). Lastly, when compared to samples treated with radiation alone, the combined treatment reduced the expression of genes overlapping with gene sets of GBM plasticity, glioma stem cells and targets of the developmental transcription factor Oct4 (**Fig. 2K/L**).

### ONC201 prolongs survival in mouse models of GBM

Based on the *in vitro* effect of ONC201 on the self-renewal capacity of patient-derived glioma specimens, we tested whether ONC201 also had anti-tumor activity in a murine model of GBM *in vivo*. In C57BL/6 mice, bearing orthotopic GL261 tumors, a single radiation dose of 10 Gy three days after implantation led to a significant increase in survival (10 Gy + Saline *v.s*. Saline: 120.5 *v.s*. 24 days, *p*<0.0001). Weekly treatment with ONC201 alone had a smaller but significant effect on median survival (ONC201 *v.s*. Saline: 42 *v.s*. 24 days, *p*<0.0001; ONC201 *v.s*. 10 Gy + Saline: 42 *v.s*. 120.5 days, *p*<0.0001). However, a single dose of 10 Gy combined with weekly ONC201 treatments led to a significant increase in survival compared to irradiation or ONC201 alone (10 Gy + ONC201 *v.s*. 10 Gy + Saline: *p*=0.0173, median survival not reached in 10 Gy + ONC201 group) with 80 % of the animals showing no signs of tumor growth 240 days after the start of the experiment (**Fig. 3A**). Importantly, ONC201 did not show any toxicity and irradiated, ONC201-treated animals showed no weight loss over the course of 240 days (**Fig. 3B**). Bioluminescence imaging of the tumors at day 3 (pre-treatment) and at day 28 (days post-implantation) showed a reduction of signal intensity after combined treatment, consistent with the observed increase in median survival (**Fig. 3C**). Likewise, while a single dose of 10 Gy led to a central necrosis of the tumors and ONC201 treatment alone had little effects, the combined treatment with radiation and ONC201 caused an almost complete regression of the tumors. (**Fig. 3D**). Staining of coronal brain sections of mice that reached euthanasia endpoints against Ki67 did not indicate a reduction in proliferation by radiation or ONC201 alone. However, the combined treatment with radiation and ONC201 decreased the number of proliferating cells at the injection site in animals euthanized at day 227 after tumor implantation (**Fig. 3E**), which was in line with the observed reduction of the bioluminescence signal (**Fig. 3C**), the reduced tumor size in H&E staining (**Fig. 3D**) and the improved median survival after combined treatment (**Fig. 3A**).

**Figure 3.**
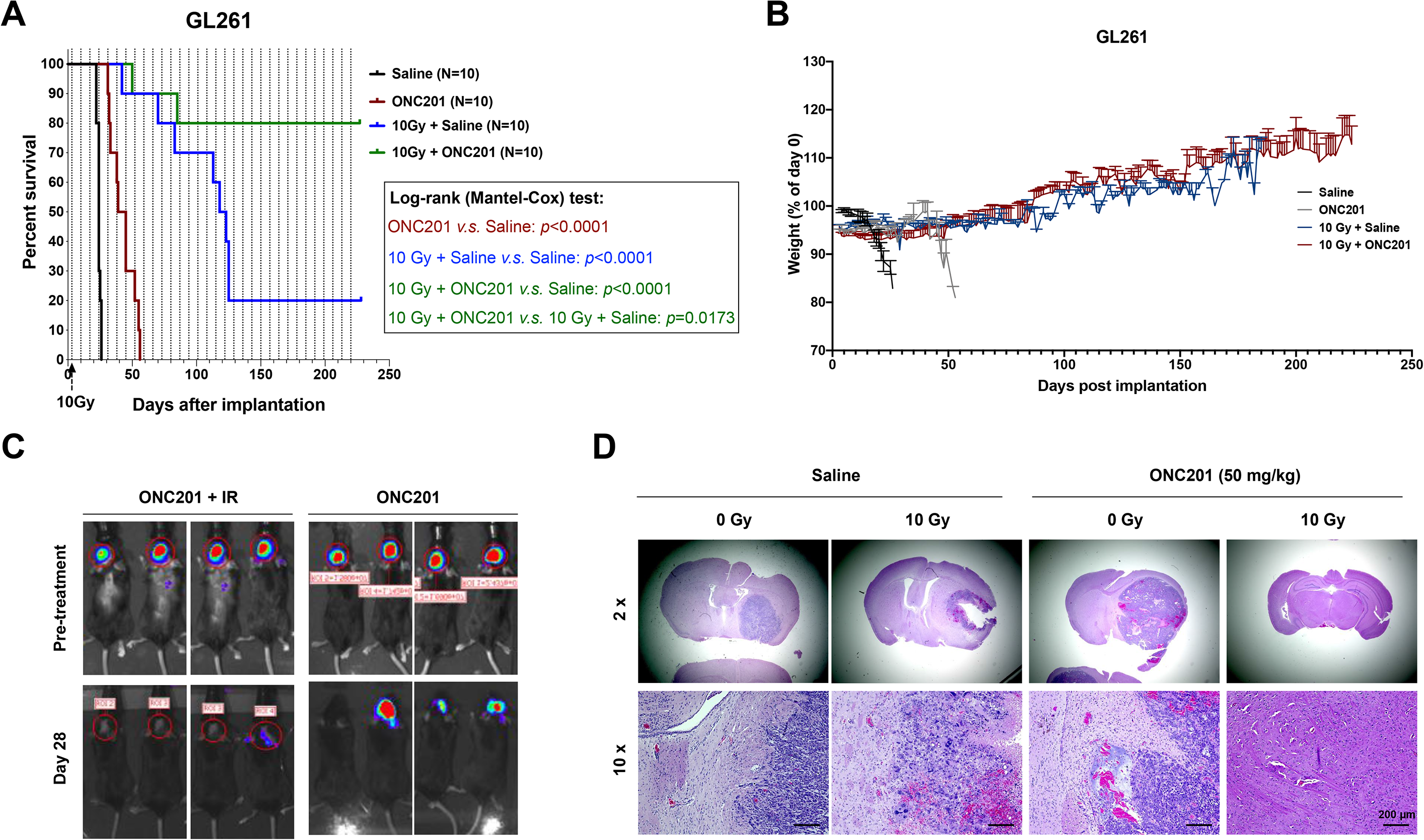
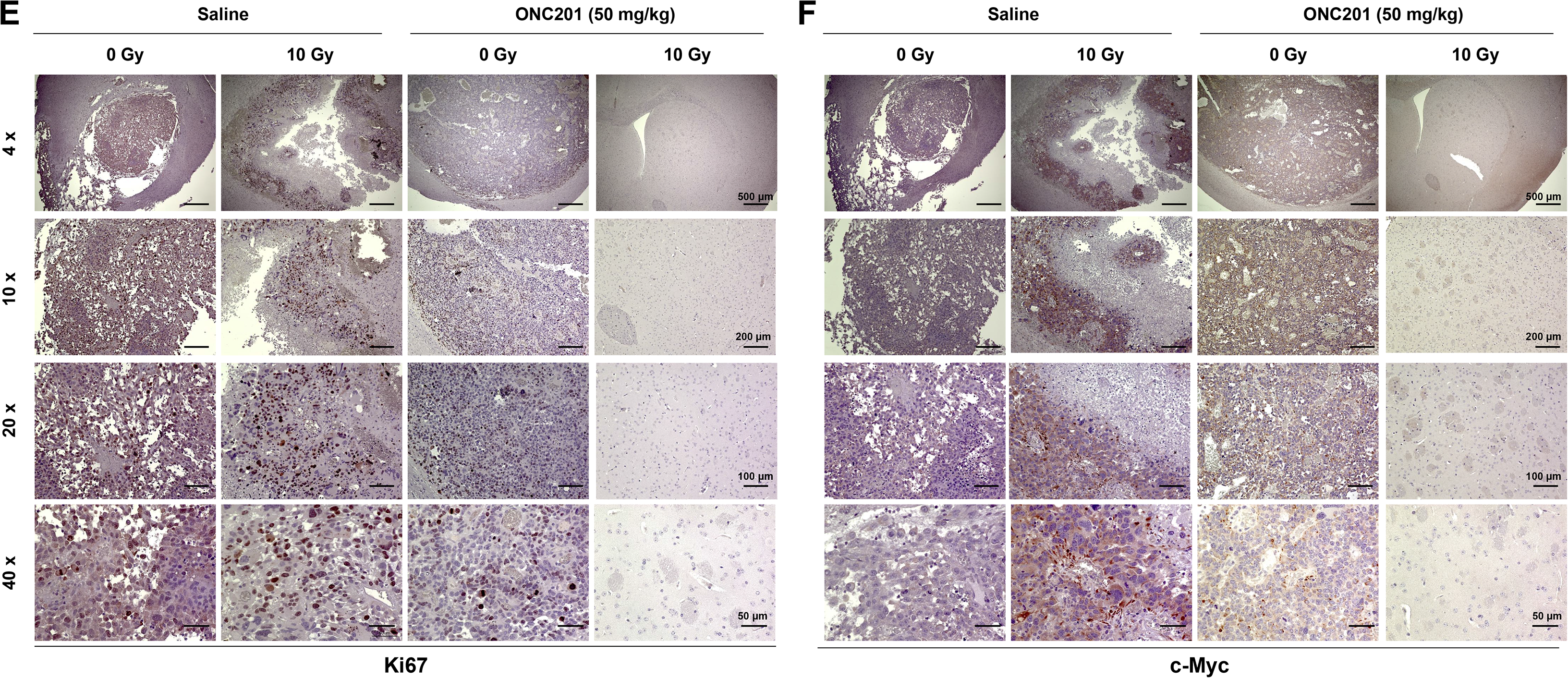
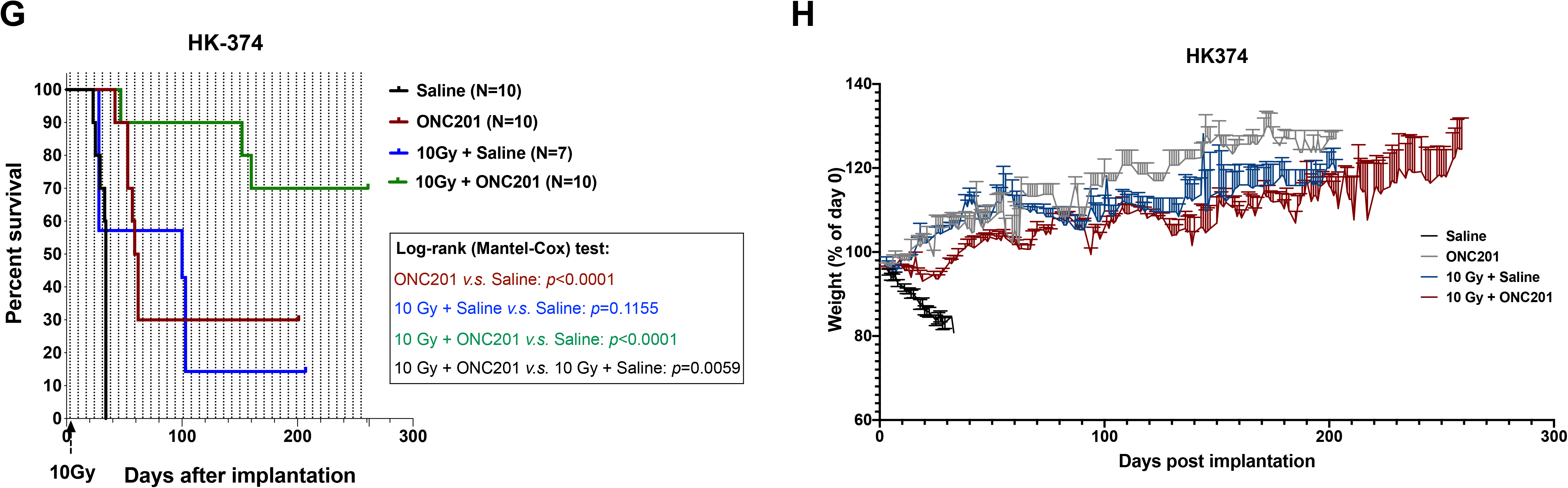
Combination of ONC201 and radiation prolongs survival in mouse models of glioblastoma. (**A**) Survival curves for C57BL/6 mice implanted intra-cranially with 2×10^5^ GL261-GFP-Luciferase mouse glioma cells. Tumors were grown for 3 days for successful grafting. Mice were irradiated with a single fraction of 0 or 10 Gy and weekly treated with Saline or ONC201 (50 mg/kg, i.p.) continuously until they reached the study endpoint. Log-rank (Mantel-Cox) test for comparison of Kaplan-Meier survival curves indicated significant differences in the ONC201 treated mice compared to their respective controls. ONC201 *v.s*. Saline (**** *p*-value<0.0001), 10 Gy + Saline *v.s*. Saline (**** *p*-value<0.0001), 10 Gy + ONC201 *v.s*. Saline (**** *p*-value<0.0001), 10 Gy + ONC201 *v.s*. 10 Gy + Saline (* *p*-value=0.0173). (**B**) Weight curves for the C57BL/6 mice in the different treatment groups. (**C**) Bioluminescence images of mice bearing tumors obtained at day 3 (pre-treatment) and at day 28 (days post implantation). (**D**) H&E stained coronal sections of the C57BL/6 mice brains implanted with GL261-GFP-Luc cells which were treated continuously with either ONC201 or saline until they met the criteria for study endpoint. (**E/F**) 4x, 10x, 20x and 40x images of Ki67 and c-Myc stained coronal sections of brains from C57BL/6 mice implanted with GL261-GFP-Luc cells and treated with either ONC201 (50 mg/kg) or saline in the presence or absence of a single dose of radiation (10 Gy) at the study endpoint. The brain samples for ONC201 in combination with radiation were from the mice at post-op day 227 (the mice did not reach the study endpoint). (**G**) Survival curves for NSG mice implanted intra-cranially with 3×10^5^ HK374-GFP-Luciferase patient-derived GBM cells. Tumors were grown for 3 days for successful grafting. Mice were irradiated with a single fraction of 0 or 10 Gy and weekly treated with Saline or ONC201 (50 mg/kg, i.p.) continuously until they reached the study endpoint. Log-rank (Mantel-Cox) test for comparison of Kaplan-Meier survival curves indicated significant differences in the ONC201 treated mice compared to their respective controls. ONC201 *v.s*. Saline (**** *p*-value<0.0001), 10 Gy + Saline *v.s*. Saline (n.s. *p*-value=0.1155), 10 Gy + ONC201 *v.s*. Saline (**** *p*-value<0.0001), 10 Gy + ONC201 *v.s*. 10 Gy + Saline (** *p*-value=0.0059). (**H**) Weight curves for the NSG mice in the different treatment groups.

Furthermore, immunohistochemistry staining revealed induction of c-Myc expression by radiation, which was in agreement with our previous report on radiation-induced acquisition of a glioma-initiating cell phenotype of previously non-tumorigenic cells^40^. This was also seen after ONC201 treatment but was completely abolished after combined treatment with radiation and ONC201 (**Fig. 3F**).

Next, we verified these findings using patient-derived orthotopic xenografts. Three days after implantation of HK-374 glioblastoma cells into the striatum of NSG mice, animals were treated with a single dose of 0 or 10 Gy followed by weekly injection of ONC201. Again, ONC201 treatment led to a small increase in median survival (ONC201 *v.s*. Saline: 60.5 *v.s*. 34 days, *p*<0.0001), while a single dose of 10 Gy alone did not significantly prolong median survival (10 Gy + Saline *v.s*. Saline: 100 *v.s*. 34 days, *p*=0.1155; 10 Gy + Saline *v.s*. ONC201: 100 *v.s*. 60.5 days, *p*=0.1472). However, the combination of a single dose of 10 Gy followed by weekly treatments with ONC201 increased overall survival when compared to 10 Gy group (10 Gy + ONC201 *v.s*. 10 Gy + Saline: *p*=0.0059, median survival not reached in 10 Gy + ONC201 group) with 70% of the animals showing no signs of tumors growth 250 days after start of the experiment (**Fig. 3G**). Similarly to what was seen in immune-competent animals, ONC201 did not show any toxicity and did not lead to weight loss in NSG mice (**Fig. 3H**).

### Altered qGBM-enriched ECM-related gene expression is associated with decreased overall survival and progression-free survival in GBM patients

To further investigate whether qGBM-enriched ECM-related gene expression is indeed associated with the clinical outcome of GBM patients, we analyzed overall survival (OS) and progression-free survival (PFS) data from 206 GBM patients in the TCGA Provisional dataset, stratified into subgroups with altered or unaltered expression of one or more ECM-related genes over-expressed in qGBM cells. Patients with increased expression of ECM-related genes showed an inferior median progression-free (5.4 *v.s*. 8.2 months, *p*-value=6.745e-3) and overall survival (11.9 *v.s*. 14.3 months, *p*-value=0.0357) compared to those with unaltered expression of these genes (**Fig. 4A/B**).

**Figure 4.**
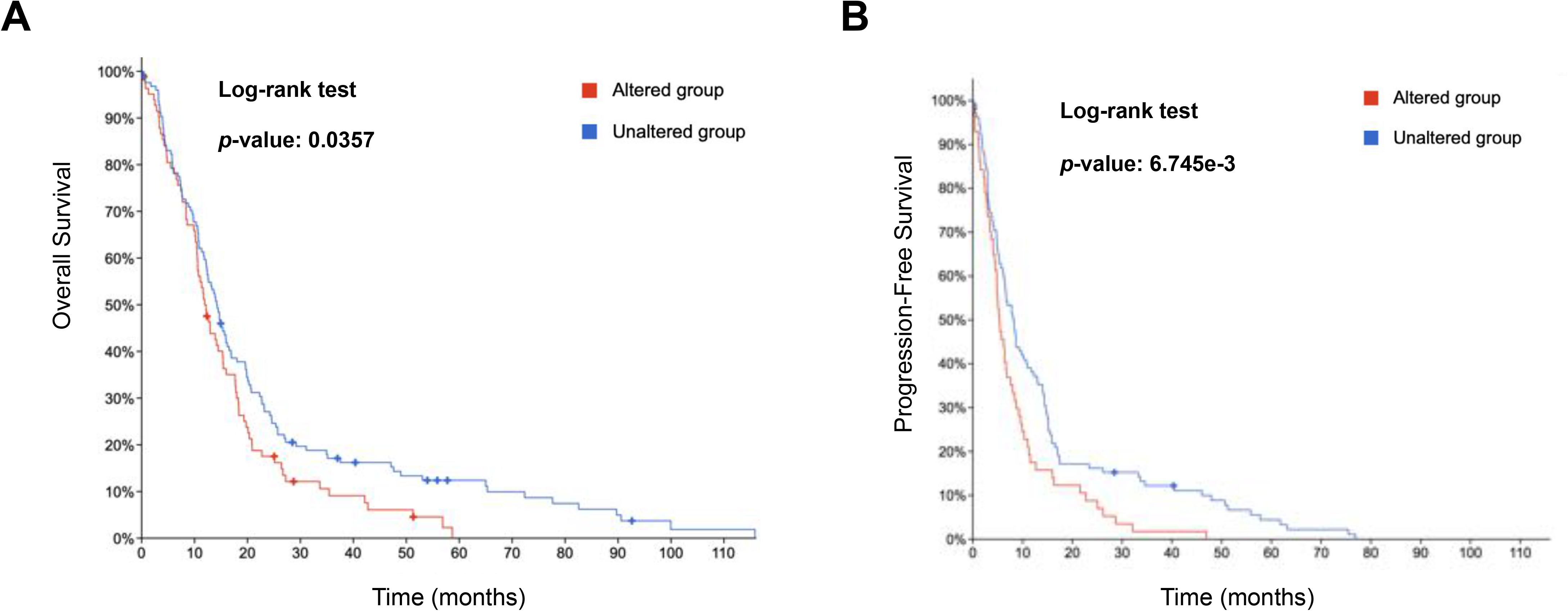
Evaluation of qGBM-enriched ECM-related gene signatures in TCGA glioblastoma patients. (**A**) The overall survival (OS) in glioblastoma patients from TCGA stratified by qGBM-enriched ECM-related gene expression. OS in qGBM-enriched ECM-related gene altered group (higher expression) was significantly worse than that of the unaltered group (log rank test, *p*-value=0.0357). (**B**) The progression-free survival (PFS) in glioblastoma patients from TCGA stratified by qGBM-enriched ECM-related gene expression. PFS in qGBM-enriched ECM-related gene altered group (higher expression) was significantly worse than that of the unaltered group (log rank test, *p*-value=6.745e-3).

## Discussion

Despite decades of research aimed at improving the treatment outcome for patients suffering from GBM, overall survival rates for this disease still remain dismal. The current standard-of-care with postoperative radiation therapy and temozolomide treatment only prolongs progression-free survival by approximately 3 months and almost all tumors recur^3^. Reasons for the failure of current therapies to control GBM are multiple and include the lack of BBB penetration of drugs used, the infiltrative nature of GBMs, intratumoral heterogeneity, and cellular plasticity^7,33,45,46^. A large number of studies support that GBM are organized hierarchically with a small number of radio- and chemotherapy-resistant glioma-initiating cells at the apex of that hierarchy, able to re-grow a tumor and give rise to differentiated progeny^47–49^.

Previous studies have shown expression of dopamine receptor 2 in GBM cells, with elevated DRD2 expression in the GIC population^50,51^. DRD2 signaling is involved in maintaining self-renewal^51^, as well as activating hypoxia response^50^ and functionally altering metabolism of GBM in a potentially autocrine manner^50^, indicating that DRD2 could be a therapeutic target in GBM.

Recent preclinical and clinical data have hinted at an efficacy of ONC201 in several solid cancers including GBM^21–23,25,27^. Originally identified as an inducer of TRAIL, ONC201 inhibits D2-like dopamine receptors DRD2 and DRD3 and activates the mitochondrial protease ClpP, a key player in the mitochondrial unfolded protein response^52,53^. So far, clinical trials using ONC201 against GBM have shown pharmacodynamic activity in biomarker-defined recurrent GBM patients, as well as in pediatric and adult H3 K27M-mutant glioma^22,26,27^.

We have recently reported that radiation in combination with the first-generation dopamine receptor antagonist trifluoperazine prolonged survival in mouse models of GBM^40^. Here we show that radiation in combination with ONC201 reduced the viability of bulk GBM cells, radiosensitized clonogenic GBM cells and reduced the self-renewal capacity of GICs *in vitro*. Combined treatment led to distinct changes in gene expression, consistent with an induction of cell death, downregulation of DNA repair and elimination of GICs. Furthermore, the combination of ONC201 and radiation prevented the radiation-induced expression of gene sets correlated with GBM plasticity, glioma stem cells, and upregulation of Oct4 targets (a developmental transcription factor^54^) and interfered with cell growth, ECM interaction and EMT genes thought to be involved in neural-to-mesenchymal transition, and the metabolism of GBM cells. This was in agreement with a previous study reporting effects of ONC201 on glioma stem cells^55^.

*In vivo*, a single dose of radiation combined with ONC201 followed by continuous, weekly ONC201 treatment translated into significantly prolonged median survival in mouse models of GBM when compared to radiation or ONC201 treatment alone. This was accompanied by tumor regression and loss of Ki67 and c-Myc staining, which was consistent with our *in vitro* data and indicated a previously unknown synergism of ONC201 and radiation. At the same time the treatment combination was well tolerated and did not lead to treatment-related toxicity.

Taken together, our data suggests that the combination of radiation with ONC201 severely affects defense mechanisms of GBM cells against radiation without acting as a classical radiosensitizer. We conclude that ONC201 in combination with radiation could be a promising new strategy against GBM that should be tested in clinical trials.

---

**Supplementary Table 1.**
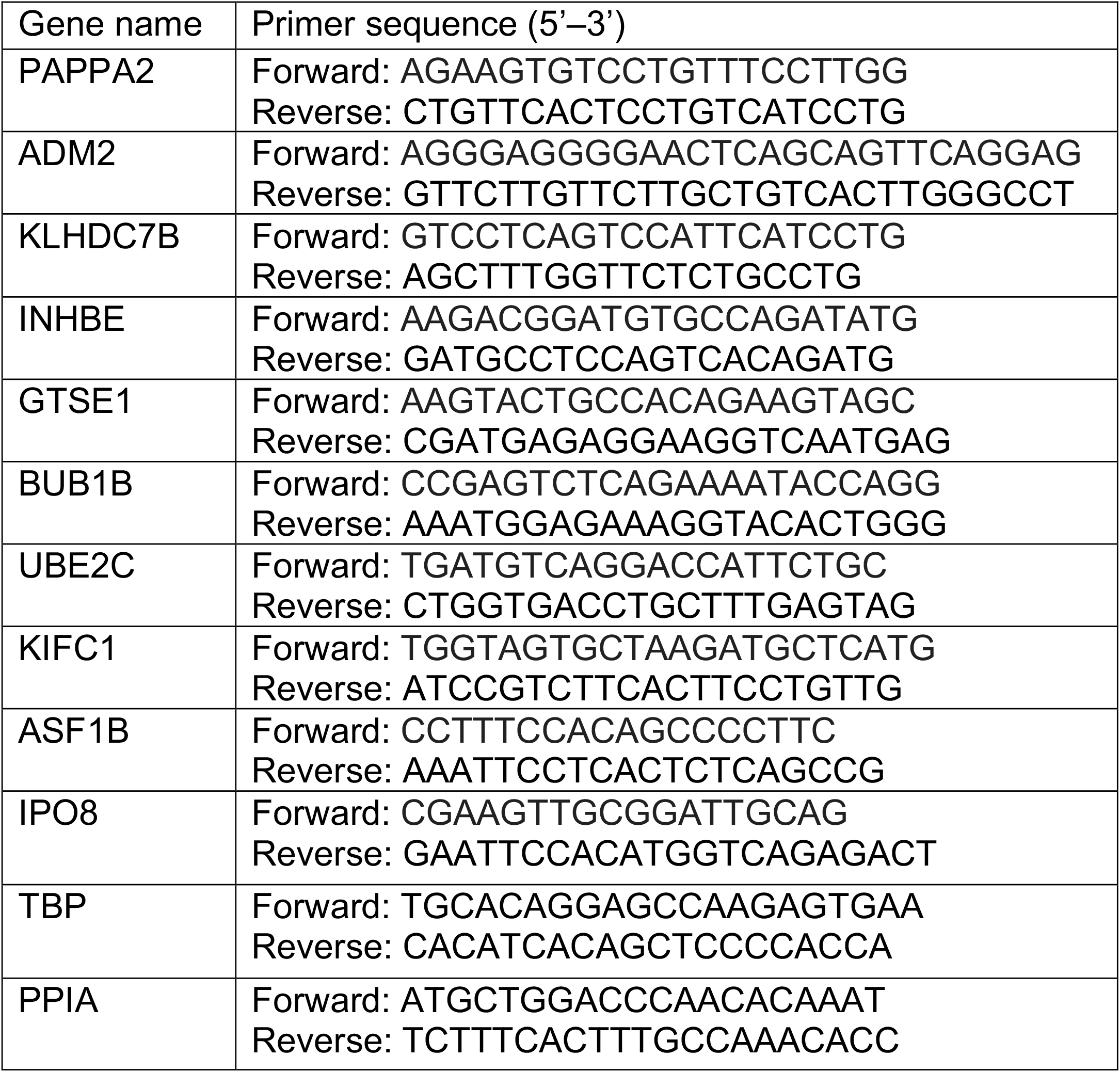

**Supplementary Figure S1.**
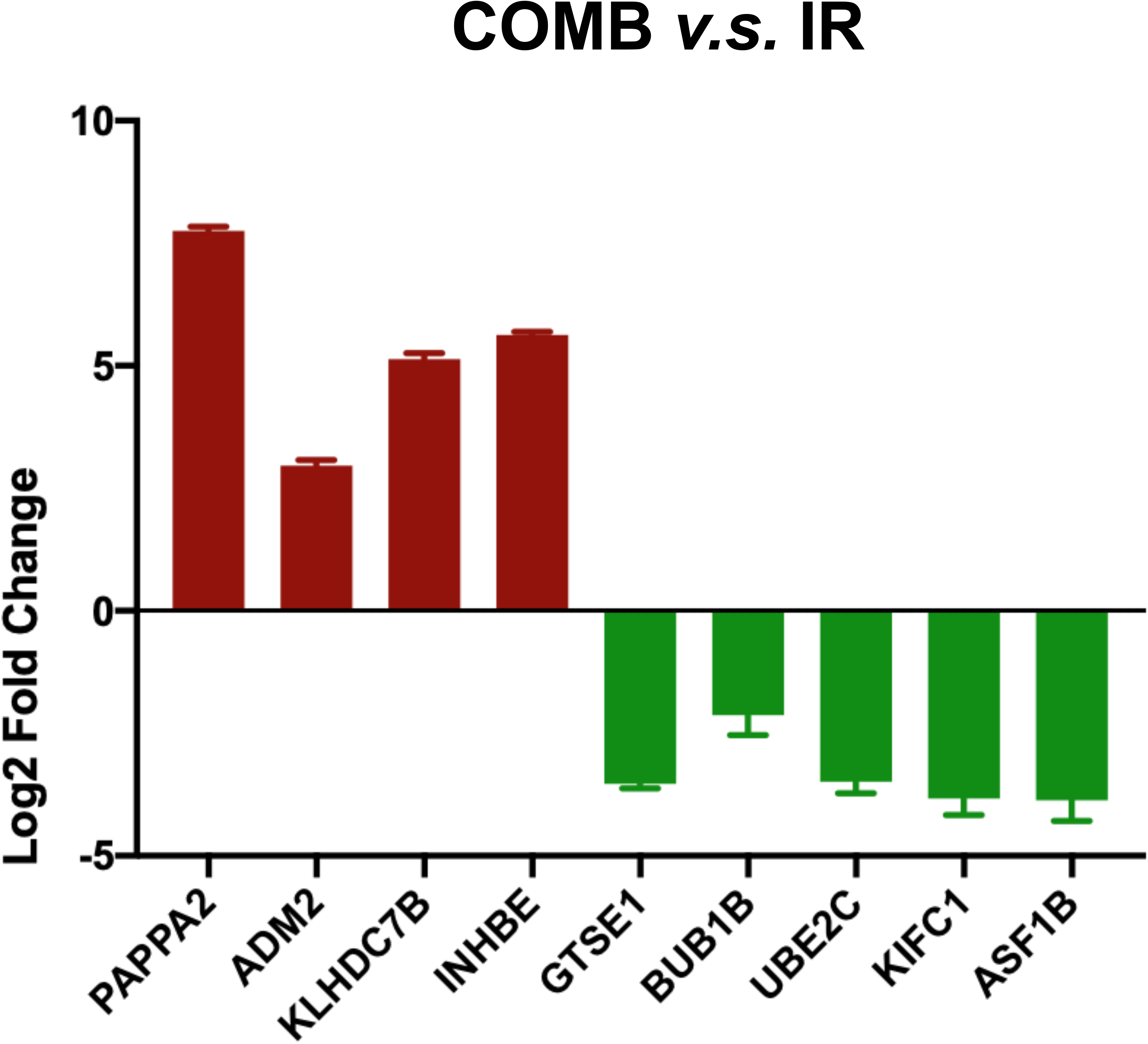
Top four up- and five down-regulated genes in cells after combined treatment with radiation and ONC201 compared to irradiated cells and their validation by qRT-PCR. All experiments have been performed with 3 biological independent repeats.

**Supplementary Figure s2.**
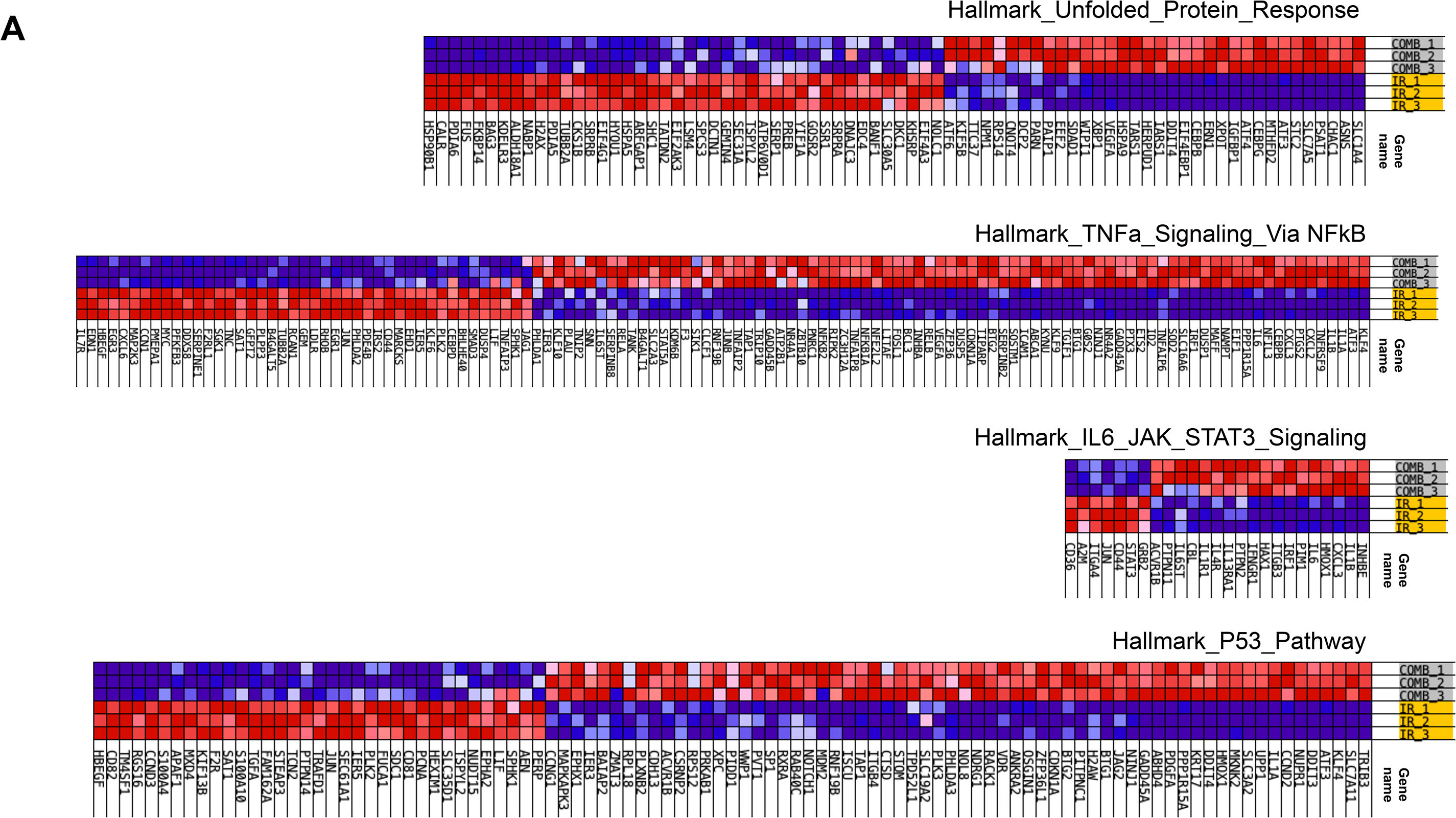
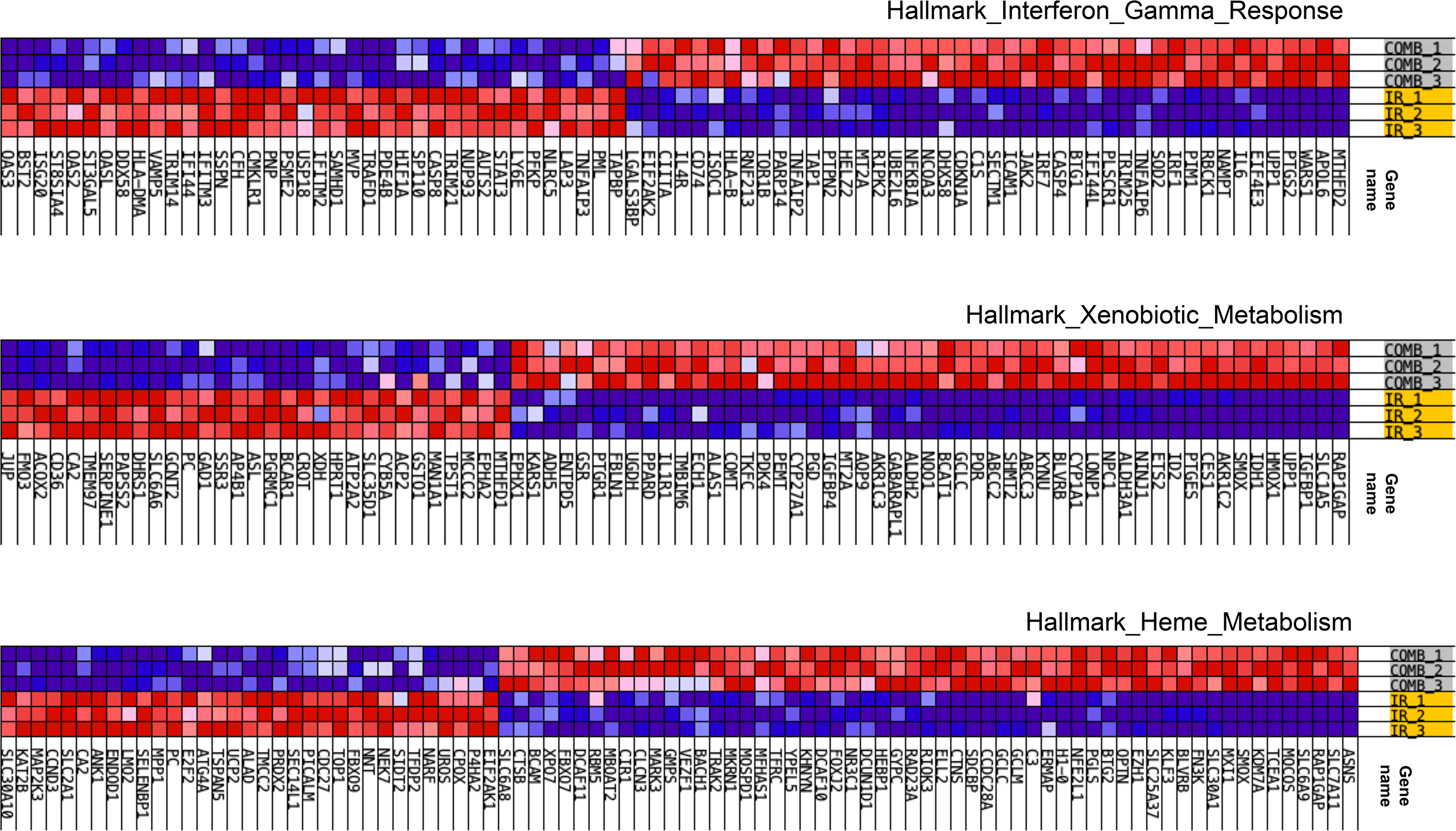
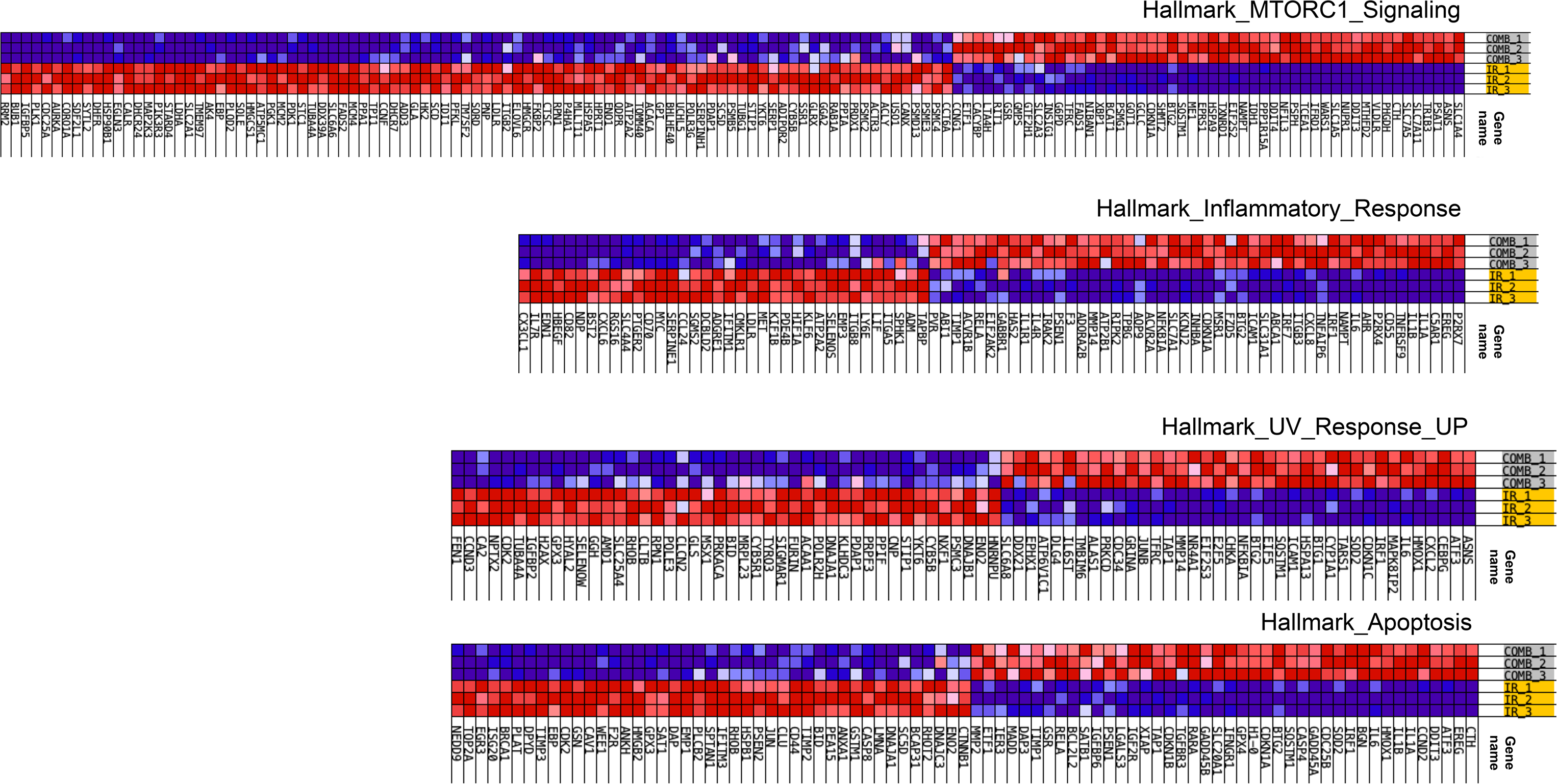
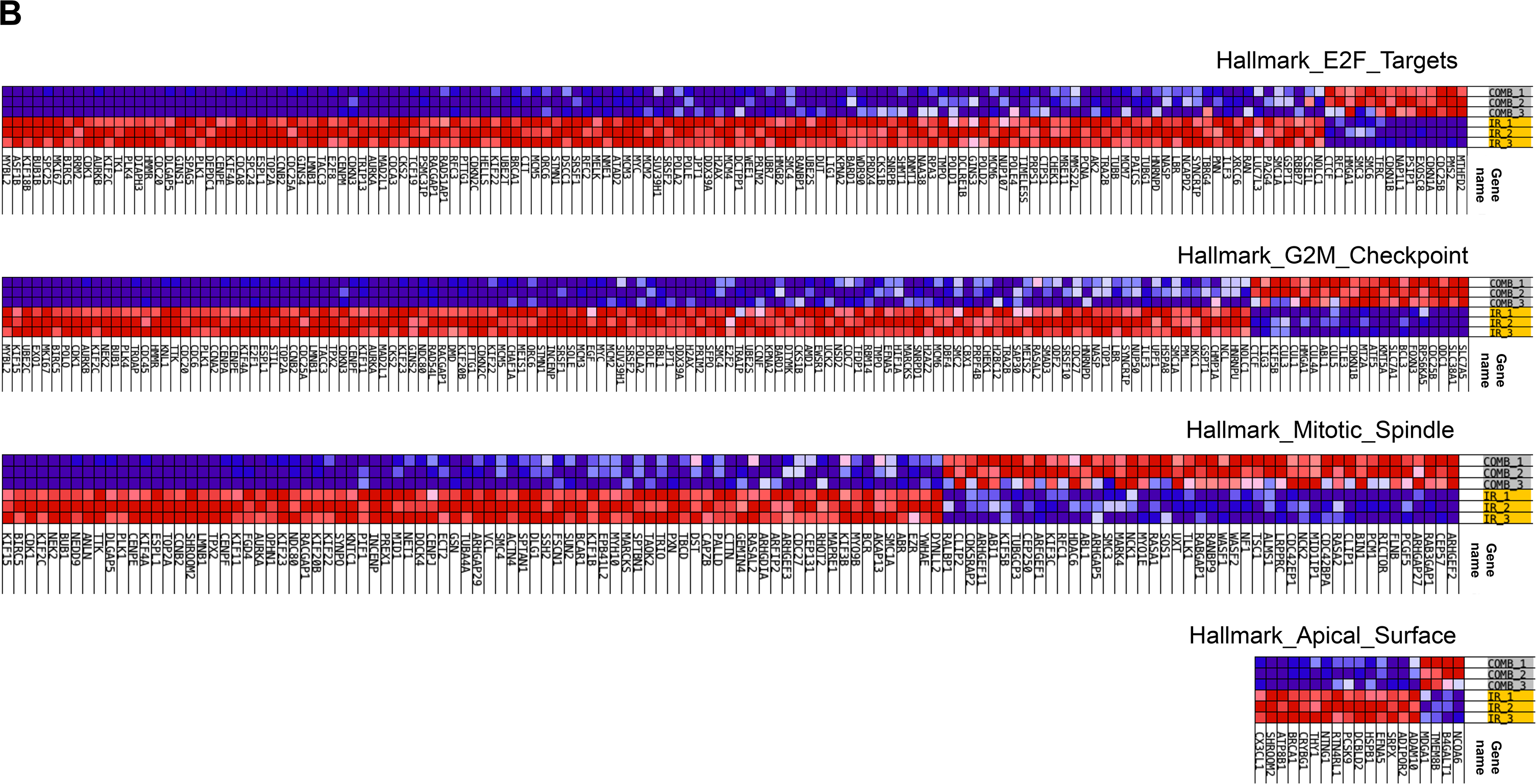
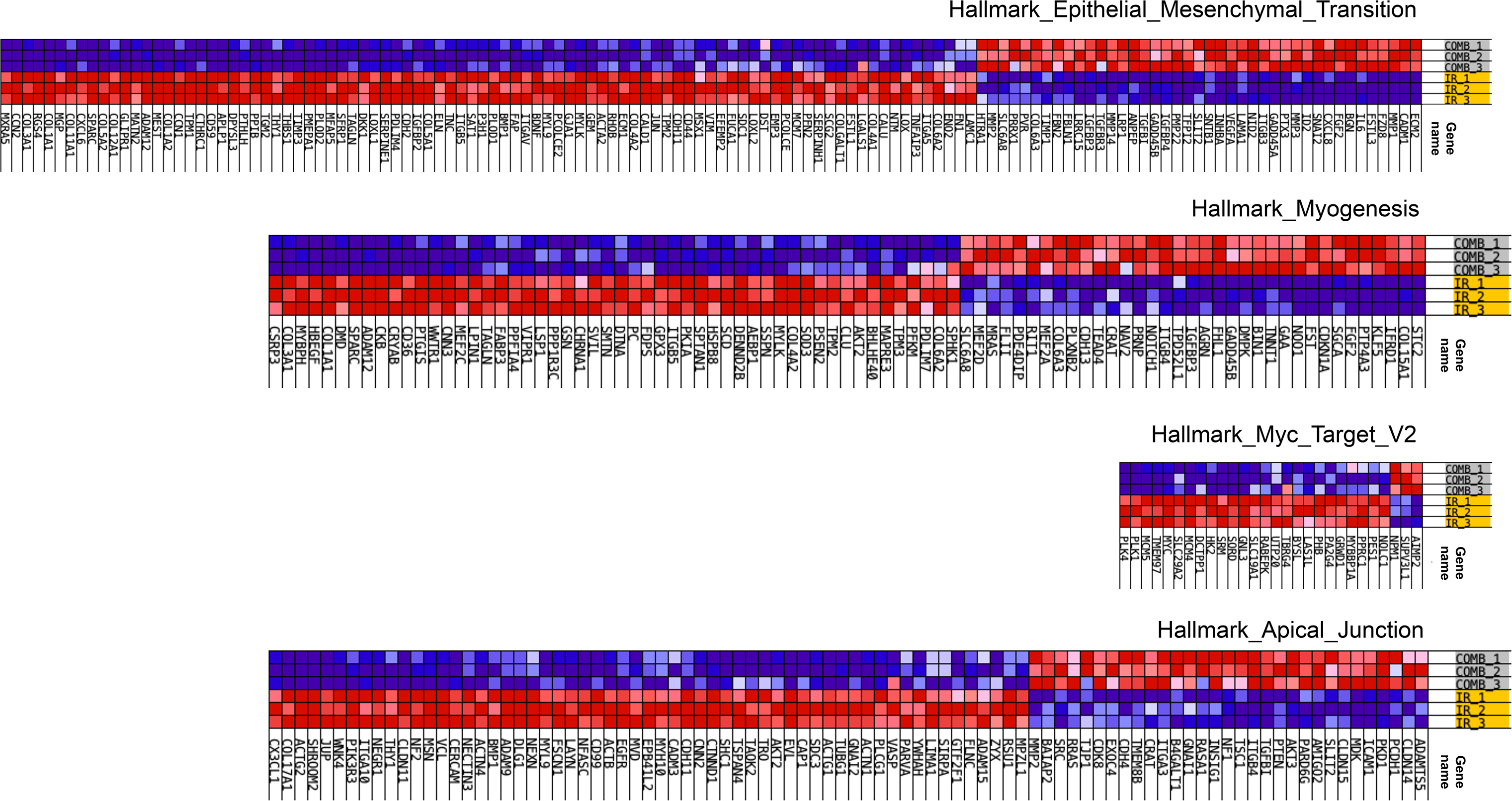
Heat maps of genes contributing to the leading edge of the up-regulated DEGs overlapped gene sets of the unfolded protein response, TNFa/NFkB, IL-6/JAK/Stat3, P53 and mTORC1 signaling, IFN gamma response, xenobiotic and heme metabolism, inflammatory, UV response and apoptosis (**A**). Heat maps of genes contributing to the leading edge of the down-regulated DEGs overlapped gene sets of E2F targets, G2/M checkpoint, Mitotic spindle, apical surface, epithelial-mesenchymal transition, myogenesis, Myc target and apical junction (**B**).

## References

1. Stupp R, Mason WP, van den Bent MJ, et al. Radiotherapy plus concomitant and adjuvant temozolomide for glioblastoma. N Engl J Med. 2005; 352(10):987–996.

2. Laperriere N, Zuraw L, Cairncross G, Cancer Care Ontario Practice Guidelines Initiative Neuro-Oncology Disease Site G. Radiotherapy for newly diagnosed malignant glioma in adults: a systematic review. Radiother Oncol. 2002; 64(3):259–273.

3. Fernandes C, Costa A, Osorio L, et al. Current Standards of Care in Glioblastoma Therapy. In: De Vleeschouwer S, ed. Glioblastoma. Brisbane (AU)2017.

4. Nicholas S, Mathios D, Ruzevick J, Jackson C, Yang I, Lim M. Current trends in glioblastoma multiforme treatment: radiation therapy and immune checkpoint inhibitors. Brain Tumor Res Treat. 2013; 1(1):2–8.

5. Chan JL, Lee SW, Fraass BA, et al. Survival and failure patterns of highgrade gliomas after three-dimensional conformal radiotherapy. J Clin Oncol. 2002; 20(6):1635–1642.

6. Osuka S, Van Meir EG. Overcoming therapeutic resistance in glioblastoma: the way forward. J Clin Invest. 2017; 127(2):415–426.

7. Dirkse A, Golebiewska A, Buder T, et al. Stem cell-associated heterogeneity in Glioblastoma results from intrinsic tumor plasticity shaped by the microenvironment. Nat Commun. 2019; 10(1):1787.

8. Auffinger B, Spencer D, Pytel P, Ahmed AU, Lesniak MS. The role of glioma stem cells in chemotherapy resistance and glioblastoma multiforme recurrence. Expert Rev Neurother. 2015; 15(7):741–752.

9. Shergalis A, Bankhead A, 3rd, Luesakul U, Muangsin N, Neamati N. Current Challenges and Opportunities in Treating Glioblastoma. Pharmacol Rev. 2018; 70(3):412–445.

10. Drean A, Goldwirt L, Verreault M, et al. Blood-brain barrier, cytotoxic chemotherapies and glioblastoma. Expert Rev Neurother. 2016; 16(11):1285–1300.

11. Laquintana V, Trapani A, Denora N, Wang F, Gallo JM, Trapani G. New strategies to deliver anticancer drugs to brain tumors. Expert Opin Drug Deliv. 2009; 6(10):1017–1032.

12. Wang D, Wang C, Wang L, Chen Y. A comprehensive review in improving delivery of small-molecule chemotherapeutic agents overcoming the blood-brain/brain tumor barriers for glioblastoma treatment. Drug Deliv. 2019; 26(1):551–565.

13. Allen JE, Krigsfeld G, Mayes PA, et al. Dual inactivation of Akt and ERK by TIC10 signals Foxo3a nuclear translocation, TRAIL gene induction, and potent antitumor effects. Sci Transl Med. 2013; 5(171):171ra117.

14. Allen JE, Krigsfeld G, Patel L, et al. Identification of TRAIL-inducing compounds highlights small molecule ONC201/TIC10 as a unique anticancer agent that activates the TRAIL pathway. Mol Cancer. 2015; 14:99.

15. Ishizawa J, Zarabi SF, Davis RE, et al. Mitochondrial ClpP-Mediated Proteolysis Induces Selective Cancer Cell Lethality. Cancer Cell. 2019; 35(5):721–737 e729.

16. Allen JE, Kline CL, Prabhu VV, et al. Discovery and clinical introduction of first-in-class imipridone ONC201. Oncotarget. 2016; 7(45):74380–74392.

17. Kline CLB, Ralff MD, Lulla AR, et al. Role of Dopamine Receptors in the Anticancer Activity of ONC201. Neoplasia. 2018; 20(1):80–91.

18. Kline CL, Van den Heuvel AP, Allen JE, Prabhu VV, Dicker DT, El-Deiry WS. ONC201 kills solid tumor cells by triggering an integrated stress response dependent on ATF4 activation by specific eIF2alpha kinases. Sci Signal. 2016; 9(415):ra18.

19. Ishizawa J, Kojima K, Chachad D, et al. ATF4 induction through an atypical integrated stress response to ONC201 triggers p53-independent apoptosis in hematological malignancies. Sci Signal. 2016; 9(415):ra17.

20. Allen JE, Prabhu VV, Talekar M, et al. Genetic and Pharmacological Screens Converge in Identifying FLIP, BCL2, and IAP Proteins as Key Regulators of Sensitivity to the TRAIL-Inducing Anticancer Agent ONC201/TIC10. Cancer Res. 2015; 75(8):1668–1674.

21. Arrillaga-Romany I, Odia Y, Prabhu VV, et al. Biological activity of weekly ONC201 in adult recurrent glioblastoma patients. Neuro Oncol. 2020; 22(1):94–102.

22. Arrillaga-Romany I, Chi AS, Allen JE, Oster W, Wen PY, Batchelor TT. A phase 2 study of the first imipridone ONC201, a selective DRD2 antagonist for oncology, administered every three weeks in recurrent glioblastoma. Oncotarget. 2017; 8(45):79298–79304.

23. Stein MN, Malhotra J, Tarapore RS, et al. Safety and enhanced immunostimulatory activity of the DRD2 antagonist ONC201 in advanced solid tumor patients with weekly oral administration. J Immunother Cancer. 2019; 7(1):136.

24. Prabhu VV, Talekar MK, Lulla AR, et al. Single agent and synergistic combinatorial efficacy of first-in-class small molecule imipridone ONC201 in hematological malignancies. Cell Cycle. 2018; 17(4):468–478.

25. Stein MN, Bertino JR, Kaufman HL, et al. First-in-Human Clinical Trial of Oral ONC201 in Patients with Refractory Solid Tumors. Clin Cancer Res. 2017; 23(15):4163–4169.

26. Hall MD, Odia Y, Allen JE, et al. First clinical experience with DRD2/3 antagonist ONC201 in H3 K27M-mutant pediatric diffuse intrinsic pontine glioma: a case report. J Neurosurg Pediatr. 2019:1–7.

27. Chi AS, Tarapore RS, Hall MD, et al. Pediatric and adult H3 K27M-mutant diffuse midline glioma treated with the selective DRD2 antagonist ONC201. J Neurooncol. 2019; 145(1):97–105.

28. Romaguera JE, Lee HJ, Tarapore R, et al. Integrated stress response and immune cell infiltration in an ibrutinib-refractory mantle cell lymphoma patient following ONC201 treatment. Br J Haematol. 2019; 185(1):133–136.

29. Hemmati HD, Nakano I, Lazareff JA, et al. Cancerous stem cells can arise from pediatric brain tumors. Proc Natl Acad Sci U S A. 2003; 100(25):15178–15183.

30. Singh SK, Hawkins C, Clarke ID, et al. Identification of human brain tumour initiating cells. Nature. 2004; 432(7015):396–401.

31. Bao S, Wu Q, McLendon RE, et al. Glioma stem cells promote radioresistance by preferential activation of the DNA damage response. Nature. 2006; 444(7120):756–760.

32. Eramo A, Ricci-Vitiani L, Zeuner A, et al. Chemotherapy resistance of glioblastoma stem cells. Cell Death Differ. 2006; 13(7):1238–1241.

33. De Vleeschouwer S, Bergers G. Glioblastoma: To Target the Tumor Cell or the Microenvironment? In: De Vleeschouwer S, ed. Glioblastoma. Brisbane (AU) 2017.

34. Matarredona ER, Pastor AM. Extracellular Vesicle-Mediated Communication between the Glioblastoma and Its Microenvironment. Cells. 2019; 9(1).

35. Tejero R, Huang Y, Katsyv I, et al. Gene signatures of quiescent glioblastoma cells reveal mesenchymal shift and interactions with niche microenvironment. EBioMedicine. 2019; 42:252–269.

36. Moore N, Lyle S. Quiescent, slow-cycling stem cell populations in cancer: a review of the evidence and discussion of significance. J Oncol. 2011; 2011.

37. Gasch C, Ffrench B, O’Leary JJ, Gallagher MF. Catching moving targets: cancer stem cell hierarchies, therapy-resistance & considerations for clinical intervention. Mol Cancer. 2017; 16(1):43.

38. Ahmed AU, Auffinger B, Lesniak MS. Understanding glioma stem cells: rationale, clinical relevance and therapeutic strategies. Expert Rev Neurother. 2013; 13(5):545–555.

39. Laks DR, Crisman TJ, Shih MY, et al. Large-scale assessment of the gliomasphere model system. Neuro Oncol. 2016; 18(10):1367–1378.

40. Bhat K, Saki M, Vlashi E, et al. The dopamine receptor antagonist trifluoperazine prevents phenotype conversion and improves survival in mouse models of glioblastoma. Proc Natl Acad Sci U S A. 2020; 117(20):11085–11096.

41. Cerami E, Gao J, Dogrusoz U, et al. The cBio cancer genomics portal: an open platform for exploring multidimensional cancer genomics data. Cancer Discov. 2012; 2(5):401–404.

42. Gao J, Aksoy BA, Dogrusoz U, et al. Integrative analysis of complex cancer genomics and clinical profiles using the cBioPortal. Sci Signal. 2013; 6(269):pl1.

43. Lathia JD, Mack SC, Mulkearns-Hubert EE, Valentim CL, Rich JN. Cancer stem cells in glioblastoma. Genes Dev. 2015; 29(12):1203–1217.

44. Pastrana E, Silva-Vargas V, Doetsch F. Eyes wide open: a critical review of sphere-formation as an assay for stem cells. Cell Stem Cell. 2011; 8(5):486–498.

45. Harder BG, Blomquist MR, Wang J, et al. Developments in Blood-Brain Barrier Penetrance and Drug Repurposing for Improved Treatment of Glioblastoma. Front Oncol. 2018; 8:462.

46. Lemee JM, Clavreul A, Menei P. Intratumoral heterogeneity in glioblastoma: don’t forget the peritumoral brain zone. Neuro Oncol. 2015; 17(10):1322–1332.

47. Goffart N, Kroonen J, Rogister B. Glioblastoma-initiating cells: relationship with neural stem cells and the micro-environment. Cancers (Basel). 2013; 5(3):1049–1071.

48. Ikushima H, Todo T, Ino Y, Takahashi M, Miyazawa K, Miyazono K. Autocrine TGF-beta signaling maintains tumorigenicity of glioma-initiating cells through Sry-related HMG-box factors. Cell Stem Cell. 2009; 5(5):504–514.

49. Niibori-Nambu A, Midorikawa U, Mizuguchi S, et al. Glioma initiating cells form a differentiation niche via the induction of extracellular matrices and integrin alphaV. PLoS One. 2013; 8(5):e59558.

50. Caragher SP, Shireman JM, Huang M, et al. Activation of Dopamine Receptor 2 Prompts Transcriptomic and Metabolic Plasticity in Glioblastoma. J Neurosci. 2019; 39(11):1982–1993.

51. Li Y, Wang W, Wang F, et al. Paired related homeobox 1 transactivates dopamine D2 receptor to maintain propagation and tumorigenicity of glioma-initiating cells. J Mol Cell Biol. 2017; 9(4):302–314.

52. Wu G, Xiong Q, Wei X, et al. Mitochondrial unfolded protein response gene CLPP changes mitochondrial dynamics and affects mitochondrial function. PeerJ. 2019; 7:e7209.

53. Qureshi MA, Haynes CM, Pellegrino MW. The mitochondrial unfolded protein response: Signaling from the powerhouse. J Biol Chem. 2017; 292(33):13500–13506.

54. Yamanaka S, Blau HM. Nuclear reprogramming to a pluripotent state by three approaches. Nature. 2010; 465(7299):704–712.

55. Prabhu VV, Lulla AR, Madhukar NS, et al. Cancer stem cell-related gene expression as a potential biomarker of response for first-in-class imipridone ONC201 in solid tumors. PLoS One. 2017; 12(8):e0180541.

